# Industrialization restructures the domestic dog gut microbiome while preserving host specificity

**DOI:** 10.64898/2026.06.29.735155

**Authors:** Arya Gautam, Dinesh Bhandari, Kusha Gurung, Aashish Gyawali, Kshitiz Gurung, Pratiksha Yadav, Kyle C. M. Smith, Anique R. Ahmad, Deena Shrestha, Kimberly Ange-van Heugten, Laura S. Weyrich, Ajit K. Karna, Aashish R Jha

## Abstract

Industrialization has reshaped human gut microbiomes, but its effects on other human-associated mammals remain poorly understood. Domestic dogs provide an informative comparative system because they have shared human environments and food systems for millennia yet retain distinct host biology. However, most canine microbiome studies have focused on industrialized companion animals, limiting our understanding of the ecological range of the domestic dog gut microbiome. We analyzed fecal 16S rRNA gene profiles from 261 dogs sampled across Nepal, Thailand, the United Arab Emirates, and the United States, spanning forager, agrarian, pastoralist, urban, and industrialized lifestyles; 257 dogs remained after excluding recent antibiotic exposure. Lifestyle was the strongest measured correlate of canine gut microbiome composition, and this structure persisted in restricted analyses of mature, non-shelter dogs sampled from temperate climate regions. Industrialized dogs differed from non-industrialized dogs through directional genus-level turnover, restructuring of VANISH- and BloSSUM-like microbial guilds, and shifts in predicted functional potential. Non-industrialized dogs were not microbiologically uniform: pastoralist dogs carried non-industrialized microbiome profiles but diverged from a simple forager-to-industrialized continuum. Cross-species comparisons with humans sampled across matched lifestyle categories showed parallel lifestyle-associated restructuring in both hosts, but host species remained the dominant axis of variation and the genera responding to industrialization were largely host-specific. These findings expand the ecological baseline for the domestic dog gut microbiome and identify industrialization as a major axis of microbiome restructuring in a long-term human-associated mammal. More broadly, they show that shared lifestyle transitions can impose parallel ecological pressures across host species without overriding host-specific community assembly.

## Introduction

Industrialization has profoundly altered the ecological conditions under which host-associated microbial communities assemble. In humans, transitions from foraging and agrarian lifestyles to urbanized and industrialized conditions are associated with changes in diet, sanitation, antibiotic exposure, environmental microbial contact, mobility, and built-environment exposure, all of which can restructure the gut microbiome [1-3]. Comparative studies across human populations have shown that industrialized populations often differ from non-industrialized populations in gut microbiome composition, prevalence of specific microbial taxa, and inferred functional potential [4-8]. These findings have led to the proposal that industrialization is associated with the loss of evolutionarily conserved bacterial lineages common in non-industrialized populations and the expansion of taxa favored in urbanized or industrialized settings [9, 10]. However, most evidence for industrialization-associated gut microbiome change comes from humans. Far less is known about whether similar ecological transitions restructure the microbiomes of other mammals that live in close association with humans.

Domestic dogs provide an informative system for addressing this gap. Dogs have lived alongside humans for thousands of years [11-13], occupying human-built and human-modified environments and consuming human-derived foods across much of their evolutionary history [14-16]. Their association with humans has tracked major transitions in subsistence, settlement, agriculture, urbanization, and industrialization and has accompanied parallel host adaptations linked to human-associated diets, including genomic changes related to starch digestion [17-21]. In contemporary industrialized settings, many dogs consume commercial diets, live in built environments, experience reduced environmental microbial exposure, and receive veterinary antibiotics and other medical interventions [22]. In contrast, free-ranging and community dogs in non-industrialized or urbanizing settings often consume household scraps, scavenged foods, livestock-associated resources, or locally available dietary substrates, while experiencing greater environmental exposure and different forms of human management. These contrasts make dogs useful for testing how lifestyle-associated ecological change shapes the gut microbiome of a long-term human-associated mammal.

Despite this potential, most canine microbiome studies have focused on industrialized companion animals, often household pets consuming commercial diets in high-income settings [23-26]. This sampling bias limits our understanding of the ecological range of the domestic dog gut microbiome and makes it difficult to determine whether the microbiome of industrialized pets represents a general canine baseline or one endpoint of a broader lifestyle-associated gradient [27, 28]. Broader sampling is especially important because dogs occupy diverse ecological roles across human societies, including free-ranging village dogs, agrarian dogs, pastoralist dogs, urban community dogs, and industrialized household pets. These populations differ in diet, environmental contact, livestock exposure, mobility, climate, geography, and human management. Characterizing this diversity is necessary for understanding how canine gut microbial communities vary across natural ecological gradients and for interpreting microbiome variation in industrialized companion animals [28].

A second unresolved question is whether lifestyle-associated microbiome change in dogs resembles changes observed in humans sharing similar ecological transitions. Across mammals and primates, gut microbiomes often retain strong host specificity, including signatures of codiversification [29, 30], host filtering [10, 31], and host-species-specific responses to human-associated environmental change [32]. At the same time, coexistence, social transmission, captivity, urbanization, and specialized diets can increase microbiome similarity across host species [33, 34]. Sympatric chimpanzees and gorillas share more gut bacterial phylotypes than allopatric hosts [34, 35], urban wildlife can acquire human-associated bacterial lineages [32], captive mammals can show partial “humanization” of the gut microbiome [36-39], and specialized diets can drive convergence across distantly related vertebrates [40]. Dogs provide a particularly useful test of this balance because they have shared human environments and food systems across generations [17, 18, 41], but remain biologically distinct hosts with their own gastrointestinal physiology, immune filtering, diet processing, and evolutionary history [42-45]. Thus, dogs can help distinguish whether shared lifestyle transitions produce taxonomic convergence across host species or instead generate parallel but host-specific microbiome responses.

To address these gaps, we analyze fecal 16S rRNA gene profiles from a globally distributed cohort of domestic dogs sampled across Nepal, Thailand, the United Arab Emirates, and the United States. The cohort spans five lifestyle categories — forager, agrarian, pastoralist, urban, and industrialized — across broad gradients of geography, altitude, climate, diet, and human association. We first test whether lifestyle is a dominant correlate of canine gut microbiome structure after accounting for major ecological and host covariates. We then identify genus-level taxa, prevalence-defined VANISH- and BloSSUM-like genera, co-abundance groups, and predicted functional profiles associated with the lifestyle gradient. Finally, we compare dogs and humans sampled across matched forager, agrarian, and industrialized contexts to test whether shared lifestyle transitions produce cross-host taxonomic convergence or host-specific microbiome responses. Because lifestyle, diet, geography, climate, and human management are inherently coupled in natural ecological systems, we treat lifestyle as an integrated ecological state rather than as a single isolated exposure. We therefore use complementary full-cohort, restricted-cohort, and projection analyses to test whether lifestyle-associated microbiome structure persists after reducing major measured sources of confounding. This framework allows us to expand the ecological baseline for the domestic dog gut microbiome while testing whether industrialization-associated microbiome restructuring extends beyond humans to another long-term human-associated mammal.

## Results

### A globally distributed canine cohort captures lifestyle transitions across geography and climate

We analyzed fecal 16S rRNA profiles from 261 domestic dogs sampled across four countries and five lifestyle categories spanning broad differences in human association, diet, management, geography, altitude, and climate (**Fig. 1A** and **Table S1**). Dogs were sampled from Nepal (n = 151), Thailand (n = 27), the United Arab Emirates (UAE; n = 13), and the United States (USA; n = 70), across altitudes ranging from 10 to 3,840 m. Sampling locations included tropical, arid-desert, temperate, and temperate-cold climate zones, providing a cohort that captured both lifestyle-associated and environmental variation.

**Figure 1.**
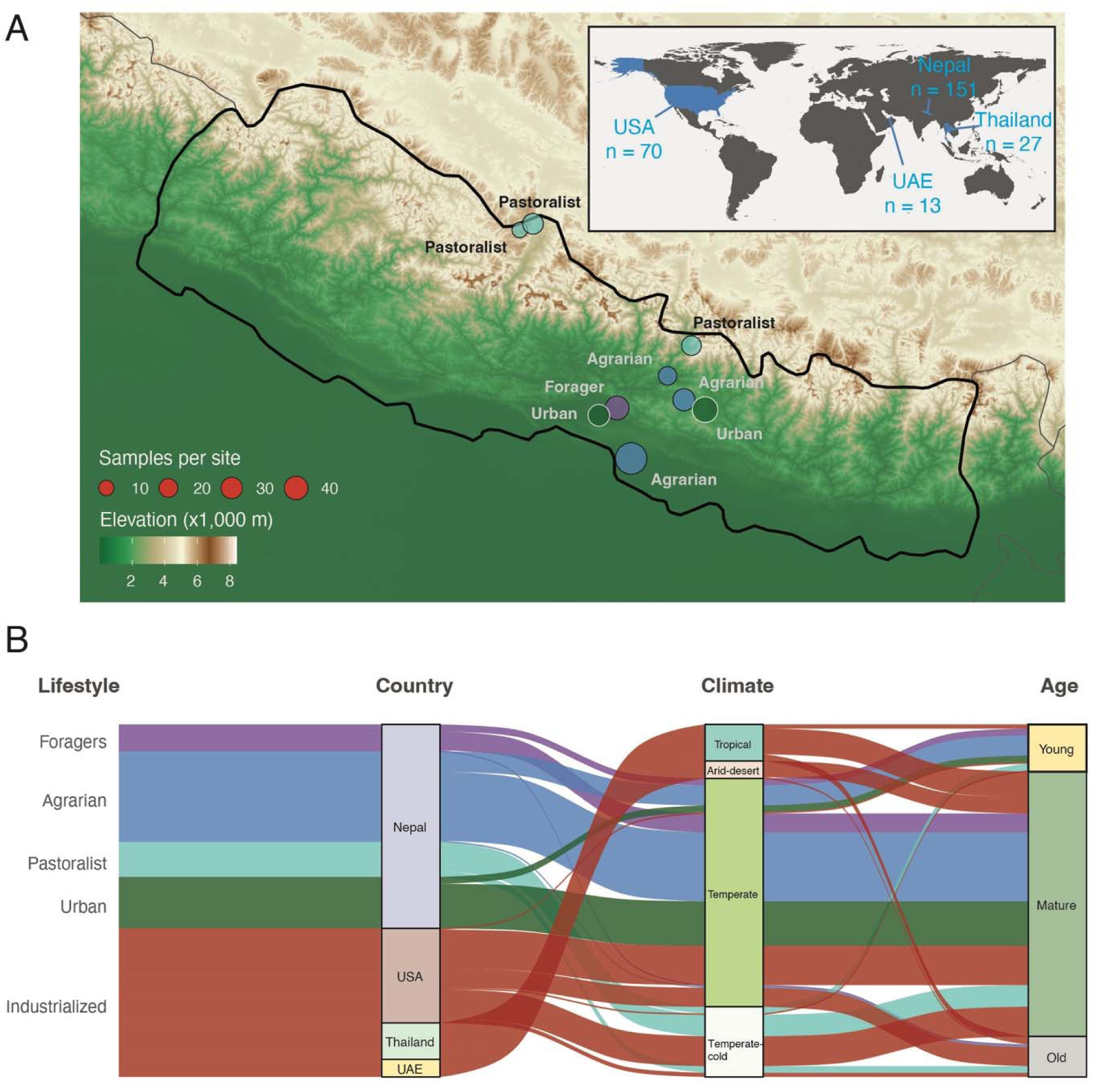
Study design and ecological structure of the globally distributed canine cohort. **(A)** Sampling locations of dogs across Nepal, Thailand, the UAE, and the United States. The main map shows Nepal sampling locations over elevation relief, with point size indicating sample number per site and labels indicating lifestyle category. The inset map shows the broader geographic distribution of sampled dogs. **(B)** Alluvial plot showing the relationship among lifestyle category, country, climate category, and age group. Flow colors indicate lifestyle categories. The figure includes all 261 fecal samples passing sequencing and contamination filtering; four dogs with reported recent antibiotic exposure were excluded from downstream microbiome analyses.

Dogs were classified into five lifestyle categories: forager, agrarian, pastoralist, urban, and industrialized. These categories reflected integrated ecological differences in human management, diet, environmental exposure, and the degree of industrialization. Forager, agrarian, pastoralist, and urban dogs were primarily free-ranging or community-associated dogs sampled in Nepal and Thailand, whereas industrialized dogs were household pets sampled in the UAE and USA. The Nepali cohort spanned four non-industrialized lifestyles across multiple communities and elevations, allowing lifestyle-associated variation to be assessed within a single country as well as across the full globally distributed cohort.

Lifestyle categories corresponded to distinct ecological contexts relevant to gut microbiome composition (**Fig. 1B**). Forager dogs were sampled from Chepang communities in Korak, where households retain sussistence links to forest-based hunting, fishing, bird trapping, wild plant gathering, and small scale maize and millet cultivation [46]. Forager and agrarian dogs primarily consumed scavenged or household-prepared foods, pastoralist dogs consumed more dairy and animal-derived foods, urban dogs had greater exposure to processed foods and urban environments, and industrialized dogs were fed predominantly commercial pet food. Environmental exposure was greatest among free-ranging dogs and lowest among household pets in industrialized settings. Thus, the cohort captures a broad ecological range of domestic dogs, from free-ranging dogs in non-industrialized communities to household pets in industrialized settings, enabling tests of how lifestyle transitions are associated with canine gut microbiome structure.

### Lifestyle is the dominant correlate of canine gut microbiome structure

After quality control, 16S rRNA gene profiling of 261 dog fecal samples generated a median of 28,617 reads per sample (mean ± SD: 33,578 ± 21,142) and identified 1,472 amplicon sequence variants (ASVs) representing 245 bacterial genera. Four dogs with recent antibiotic exposure showed reduced diversity and were excluded, leaving 257 dogs for downstream analyses.

Principal coordinate analysis (PCoA) of genus-level weighted UniFrac distances showed strong structuring of gut microbiome composition by lifestyle (**Fig. 2A**). The first two PCoA axes explained 42.9% of the variation, with PCo1 explaining 29.9% and separating many industrialized dogs from non-industrialized dogs. PCo2 explained an additional 13.0% and captured secondary differentiation among non-industrialized groups, with pastoralist dogs deviating from a simple forager-to-industrialized gradient. Notably, dogs sampled in Thailand clustered closer to dogs from the UAE and United States than to most non-industrialized Nepali dogs (**Fig. S1)**, consistent with their sampling from an urbanizing food environment in Chiang Mai [47].

**Figure 2.**
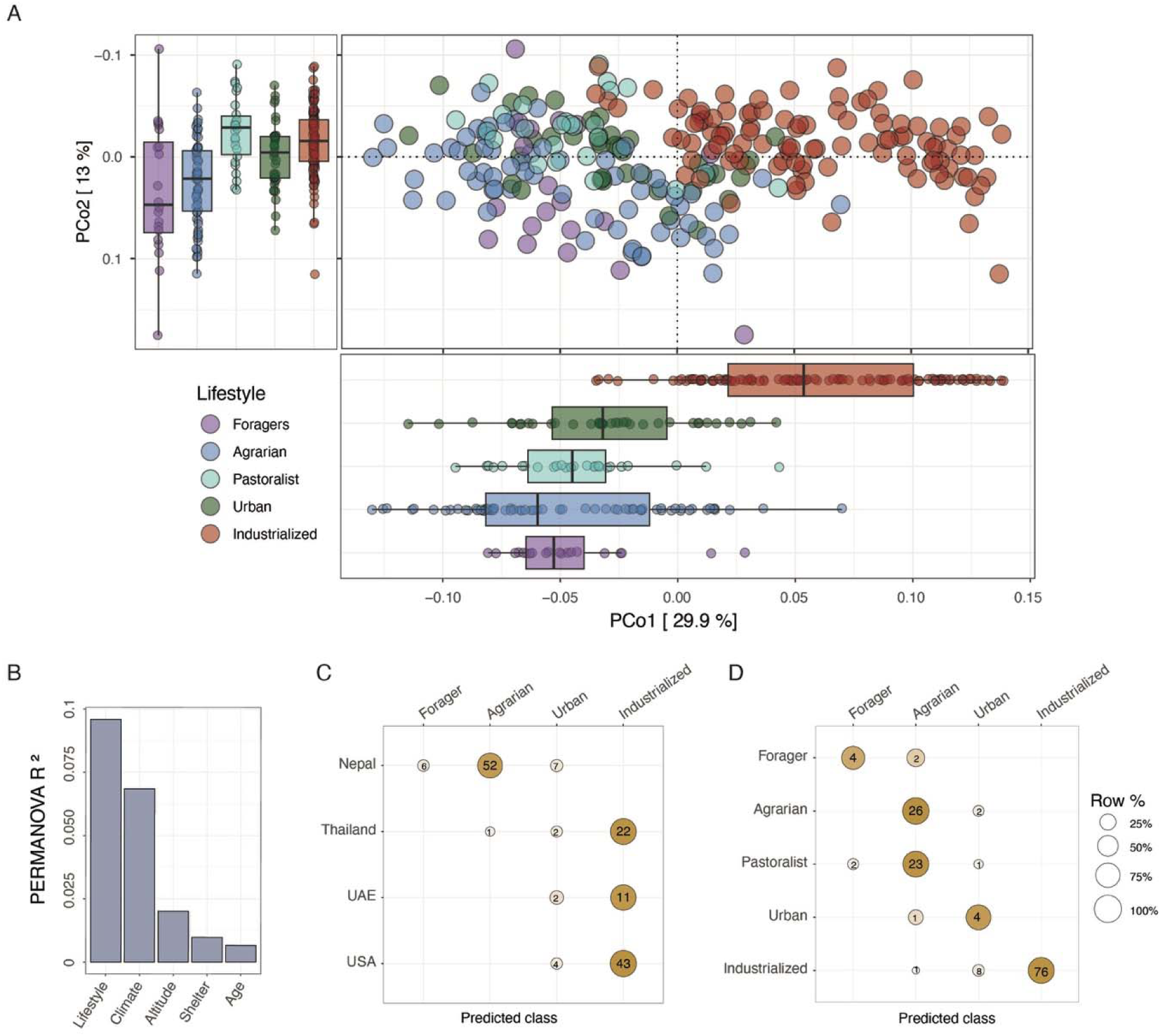
Lifestyle-associated structure in the canine gut microbiome. **(A)** Weighted UniFrac PCoA of genus-level microbiome profiles from 257 dogs. Points represent individual dogs and are colored by lifestyle. Marginal boxplots show sample distribution along PCo1 and PCo2 by lifestyle. **(B)** Marginal PERMANOVA R² values from a multivariable model including lifestyle, climate category, altitude category, shelter status, and age group. **(C)** Projection of dogs excluded from the restricted random forest training set, summarized by country. **(D**) Projection of the same excluded dogs summarized by observed lifestyle category. The classifier was trained on mature, non-shelter dogs from temperate regions, excluding Thai and UAE dogs. Balloon size indicates the row-wise percentage of samples assigned to each predicted class, and numbers indicate sample counts. Zero-count cells are omitted.

We quantified associations between microbiome structure and measured host or ecological variables using complementary approaches. Marginal PERMANOVA tested associations with the full genus-level distance matrix, whereas EnvFit tested alignment with the leading ordination axes. In PERMANOVA models using weighted UniFrac distances, lifestyle explained the largest share of overall microbiome variation among measured variables (R² = 0.10, P = 0.001), followed by climate category (R² = 0.07, P = 0.001). Altitude category explained a smaller fraction of variation (R² = 0.02, P = 0.001), as did shelter status (R² = 0.01, P = 0.003), whereas age group was not significant (R² = 0.007, P = 0.186; **Fig. 2B, Table S2**). Results were consistent using Bray-Curtis distances (**Fig. S2, Table S2**).

EnvFit analyses using the first three ordination axes supported the same overall pattern (**Table S2**). Lifestyle showed the strongest alignment with ordination structure (R² = 0.43 and 0.49, P = 0.001 for weighted UniFrac and Bray-Curtis, respectively) followed by climate category (R² = 0.27 for both distance metrics, P = 0.001) and altitude group (R² = 0.22 and 0.26, P = 0.001). Shelter status was not significant in EnvFit analyses (P > 0.05). Although age group explained little variation overall, a small association was detected in EnvFit for both weighted UniFrac and Bray-Curtis ordinations (R² = 0.037 and 0.038, P = 0.001). Thus, lifestyle was the strongest measured correlate of canine gut microbiome composition across both distance-based and ordination-based analyses, with climate contributing additional structure.

Because lifestyle, climate, age, shelter status, and geography are partially coupled in this observational cohort, we next tested whether lifestyle-associated structure persisted after reducing major measured sources of confounding. We repeated beta-diversity analyses in a restricted subset of mature, non-shelter dogs sampled from temperate climate regions, excluding Thai and UAE dogs. This subset retained 107 dogs spanning forager, agrarian, urban, and industrialized lifestyles. Lifestyle remained strongly associated with microbiome composition in both weighted UniFrac and Bray-Curtis analyses, explaining nearly 20% of variation in each distance metric (weighted UniFrac: R² = 0.197, P = 0.001; Bray-Curtis: R² = 0.196, P = 0.001; PERMANOVA, **Table S2, Fig. S3**). Restricted EnvFit showed similarly strong alignment between lifestyle and ordination structure (weighted UniFrac: R² = 0.33, P = 0.001; Bray-Curtis: R² = 0.35, P = 0.001; **Table S2**). These results indicate that lifestyle-associated microbiome structure was not explained solely by measured differences in climate, age, shelter status, or inclusion of Thai and UAE dogs.

We then asked whether lifestyle-associated microbiome states learned from this restricted dataset could classify dogs excluded from the restricted training set. A random forest classifier trained on the restricted mature, non-shelter, temperate-climate subset recovered lifestyle categories with high held-out accuracy (accuracy = 0.864, Cohen’s κ = 0.806; **Fig. S4A**), comparable to the classifier trained on the full dataset with 257 dogs (**Fig. S4B**). When excluded dogs were projected onto this classifier, assignments were biologically coherent by both country and lifestyle (**Fig. 2C, D**). No excluded Nepali dogs were classified as industrialized; instead, most were assigned to agrarian, forager, or urban classes. In contrast, most Thai, UAE, and US dogs were assigned to the industrialized class, including 22 of 25 Thai urban community dogs, 11 of 13 UAE dogs, and 43 of 47 excluded US dogs. The remaining UAE and US dogs were assigned to the urban class rather than to agrarian or forager classes. These projections supported the placement of Thai urban community dogs near the urbanized/industrialized end of the microbiome gradient, while retaining their ecological interpretation as urban community dogs rather than industrialized household pets.

The projection analysis also clarified the position of pastoralist dogs. Because pastoralist dogs were not included as a training class in the restricted classifier, they were assigned to the closest available lifestyle category. Most pastoralist dogs were classified as agrarian, with a small number assigned to forager or urban classes and none assigned to industrialized. This indicates that pastoralist dogs carried non-industrialized microbiome profiles, while also supporting their treatment as a distinct ecological group that does not fall neatly along a simple forager-to-industrialized continuum.

Although altitude aligned with the leading ordination axes in the full dataset, its marginal contribution in PERMANOVA was small. This likely reflected the position of high-altitude pastoralist dogs along PCo2, because all high-altitude dogs in this cohort were pastoralist dogs. We therefore interpreted altitude as part of the integrated pastoralist ecological context rather than as an independent predictor in downstream analyses. Downstream models retained lifestyle and climate as primary ecological predictors and age group as a conservative host covariate; shelter status was not retained because it explained little variation overall and applied to a limited subset of dogs.

To assess whether the lifestyle signal was driven primarily by broad international contrasts, we repeated the ordination, PERMANOVA, and EnvFit analyses within Nepal. In both distance metrics, Nepali dogs showed lifestyle-associated structure within a single country, with pastoralist dogs again departing from the simple forager-to-urban continuum (weighted UniFrac: R² = 0.08, P = 0.001; Bray-Curtis: R² = 0.09, P = 0.001; **Fig. S5**). Importantly, lifestyle-associated microbiome structure was also detectable within Nepal alone, indicating that the signal was not reducible to broad country-level differences.

Finally, we asked whether within-sample diversity followed the same ecological pattern observed for community composition. In generalized linear models including lifestyle and other host and ecological covariates, alpha diversity showed no significant associations with measured predictors after Benjamini-Hochberg correction across all alpha diversity tests, whether measured as observed ASV richness, Shannon diversity, or Faith’s phylogenetic diversity (**Table S3**). Thus, lifestyle-associated variation among dogs was reflected primarily in microbiome structure rather than in overall within-sample diversity.

### Lifestyle is associated with directional restructuring of the canine gut microbiota

To identify bacterial genera underlying the compositional gradient, we performed genus-level differential abundance analysis using MaAsLin2 [48]. In a lifestyle-only model with industrialized dogs as the reference group, 142 of 245 genera showed significant large-effect differences in at least one non-industrialized lifestyle group relative to industrialized dogs (FDR-adjusted q < 0.05 and an absolute MaAsLin2 coefficient > 1; **Table S4**). This broad signal indicated extensive genus-level restructuring across canine lifestyle categories.

To focus on taxa contributing to the main lifestyle gradient, we next identified genera showing monotonic abundance patterns across the forager-agrarian-urban-industrialized axis. Pastoralist dogs were excluded from this monotonic-gradient definition because they represented a distinct high-altitude lifestyle rather than a simple intermediate state along the forager-to-industrialized continuum. This analysis identified 48 genera with significant large endpoint contrasts between forager and industrialized dogs (FDR-adjusted q < 0.05 and absolute coefficient > 1; **Table S5**). These genera separated into two opposing patterns: genera enriched in forager dogs and reduced toward industrialized dogs, and genera showing the reverse pattern (**Fig. 3A**).

**Figure 3.**
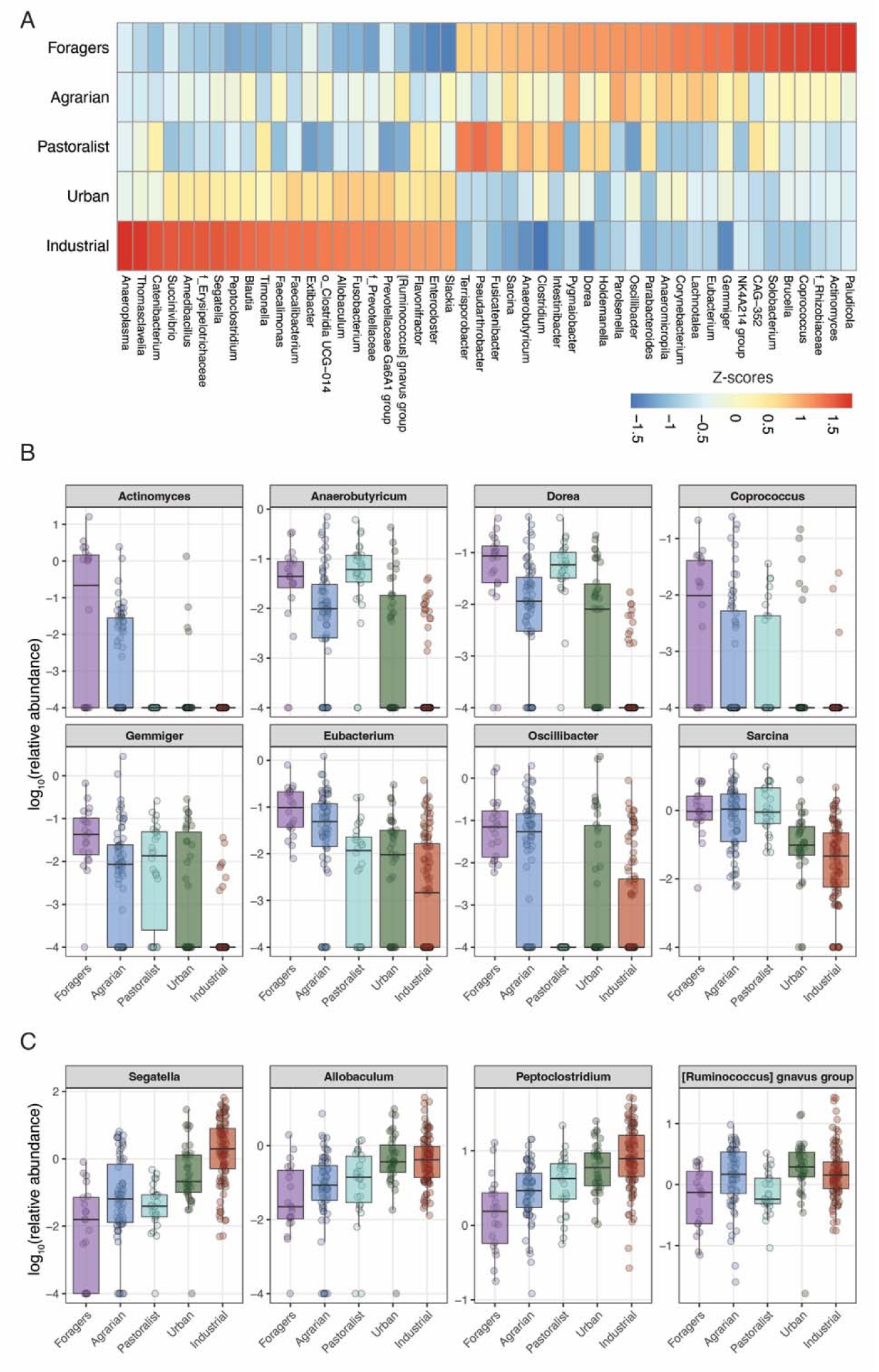
Directional genus-level shifts across the canine lifestyle gradient. **(A)** Heatmap of genera showing monotonic lifestyle-associated abundance patterns across the forager-agrarian-urban-industrialized axis. Values show row-scaled mean relative abundance within each genus. Red indicates higher relative abundance and blue indicates lower relative abundance within each genus. Genera were retained if they showed significant and large endpoint contrasts between forager and industrialized dogs (FDR-adjusted q < 0.05 and absolute MaAsLin2 coefficient > 1). **(B)** Representative genera enriched in non-industrialized dogs and declining toward industrialized dogs. **(C)** Representative genera enriched in industrialized dogs and reduced in forager or agrarian dogs. Boxplots in **B** and **C** show log-transformed relative abundance across lifestyle groups, with points representing individual dogs.

Genera enriched in non-industrialized dogs included several anaerobic gut lineages previously associated with carbohydrate fermentation or short-chain fatty acid production, including *Actinomyces*, *Anaerobutyricum*, *Dorea*, *Sarcina*, *Coprococcus*, *Gemmiger*, *Eubacterium*, and *Oscillibacter* (**Fig. 3B)** [49-54]. Additional genera enriched in non-industrialized dogs, including *Corynebacterium* and *Brucella* may reflect broader environmental or livestock-associated exposures [55, 56]. These patterns are consistent with, but do not directly demonstrate, differences in dietary substrate use and environmental exposure across lifestyles.

Industrialized dogs showed higher relative abundance of a distinct set of genera, including Prevotellaceae-associated taxa such as *Segatella*, Erysipelotrichaceae-associated genera such as *Allobaculum* and *Catenibacterium*, and anaerobic gut lineages including *Blautia*, *Enterocloster*, *Flavonifractor*, *Peptoclostridium*, and the [Ruminococcus] gnavus group (**Fig. 3C)**. Thus, industrialization in dogs was associated not only with reduced abundance of several non-industrialized-associated anaerobes, but also with enrichment of distinct gut bacterial genera.

To determine whether these lifestyle-associated shifts were explained by covarying ecological or host variables, we fit a multivariable model including lifestyle, climate category, and age group. Of the 142 large-effect lifestyle-associated genera identified in the univariate model, 137 remained significant after adjustment (FDR-adjusted q < 0.05 and an absolute MaAsLin2 coefficient > 1; **Table S6**). Across shared genus-lifestyle contrasts, effect directions were fully concordant between the lifestyle-only and multivariable models, and coefficients were strongly correlated before and after adjustment (r = 0.98; **Fig. S6**). The median absolute coefficient changed only modestly, from 2.19 to 2.10. These results indicate that lifestyle-associated genus-level shifts were largely preserved after accounting for climate and age.

### Climate contributes additional, zone-specific genus-level structure

Because climate was consistently associated with microbiome composition in beta-diversity analyses, we next examined genus-level climate associations in the multivariable model that included lifestyle, climate category, and age group. Eighty-three genera showed significant large-effect associations with climate category using tropical as the reference (FDR-adjusted q < 0.05 and an absolute MaAsLin2 coefficient > 1; **Table S6**). Lifestyle and climate signals partially overlapped: 66 genera were associated with both variables, whereas 71 lifestyle-associated genera were not climate-associated, indicating that lifestyle captured substantial genus-level variation beyond broad climatic differences (**Fig. S7**).

Climate-associated genera showed a different pattern from lifestyle-associated genera. Whereas lifestyle-associated genera formed a directional forager-to-industrialized gradient, climate-associated genera formed more discrete climate-enriched blocks (**Fig. S8**). After excluding genera that were also associated with lifestyle or age, 17 climate-specific genera remained. Most were enriched in tropical environments and included environment-associated genera such as *Sphingobacterium*, *Chryseobacterium*, and *Empedobacter* (**Fig. S8**), suggesting that warm, humid environments may contribute exposure-related signals not captured by lifestyle alone.

Climate-associated differences were detectable within industrialized dogs sampled across tropical, arid-desert, temperate, and temperate-cold climate categories (P < 0.05, PERMANOVA; **Fig. S9**). This supported climate and geography as additional sources of structured variation, although these effects were weaker than the primary lifestyle-associated gradient. In contrast, age-related genus-level associations were limited: only nine genera showed significant large-effect associations with age after accounting for lifestyle and climate, including a decline in *Bifidobacterium* with age (**Fig. S10; Table S6**) [57]. Thus, climate contributed additional zone-specific microbiome structure, whereas age had comparatively modest effects in this cohort.

### Industrialization is associated with coordinated turnover of VANISH-like and BloSSUM-like genera in dogs

We next asked whether lifestyle-associated taxonomic shifts in dogs resembled prevalence-based patterns previously linked to human industrialization. In humans, VANISH taxa are more prevalent in non-industrialized populations, whereas BloSSUM taxa are more prevalent in urbanized or industrialized populations [4, 58]. We applied this prevalence-based framework to dogs by comparing forager and agrarian dogs with dogs from industrialized settings.

This analysis identified 61 genera with large prevalence differences between non-industrialized and industrialized dogs, including 46 VANISH-like and 15 BloSSUM-like genera (absolute prevalence difference ≥ 30%, FDR-adjusted P < 0.05, *Fisher’s exact test*; **Fig. 4A** and **Table S7**). VANISH-like genera included multiple anaerobic gut lineages, including *Coprococcus, Dorea, Ruminococcus*, *Oscillibacter*, *Anaerobutyricum*, *Anaeromicropila*, *Eubacterium*, *Intestinibacter*, and Christensenellaceae-associated taxa. BloSSUM-like genera included *Flavonifractor, Timonella, Tyzzerella, Anaeroplasma, Absiella, Empedobacter*, and *Streptococcus*, and several family-level assignments within Erysipelotrichaceae, Muribaculaceae, Enterococcaceae, and Peptostreptococcaceae. Thus, industrialized dogs showed both reduced prevalence of a broad set of non-industrialized-associated anaerobes and increased prevalence of a smaller set of industrialized-associated taxa.

**Figure 4.**
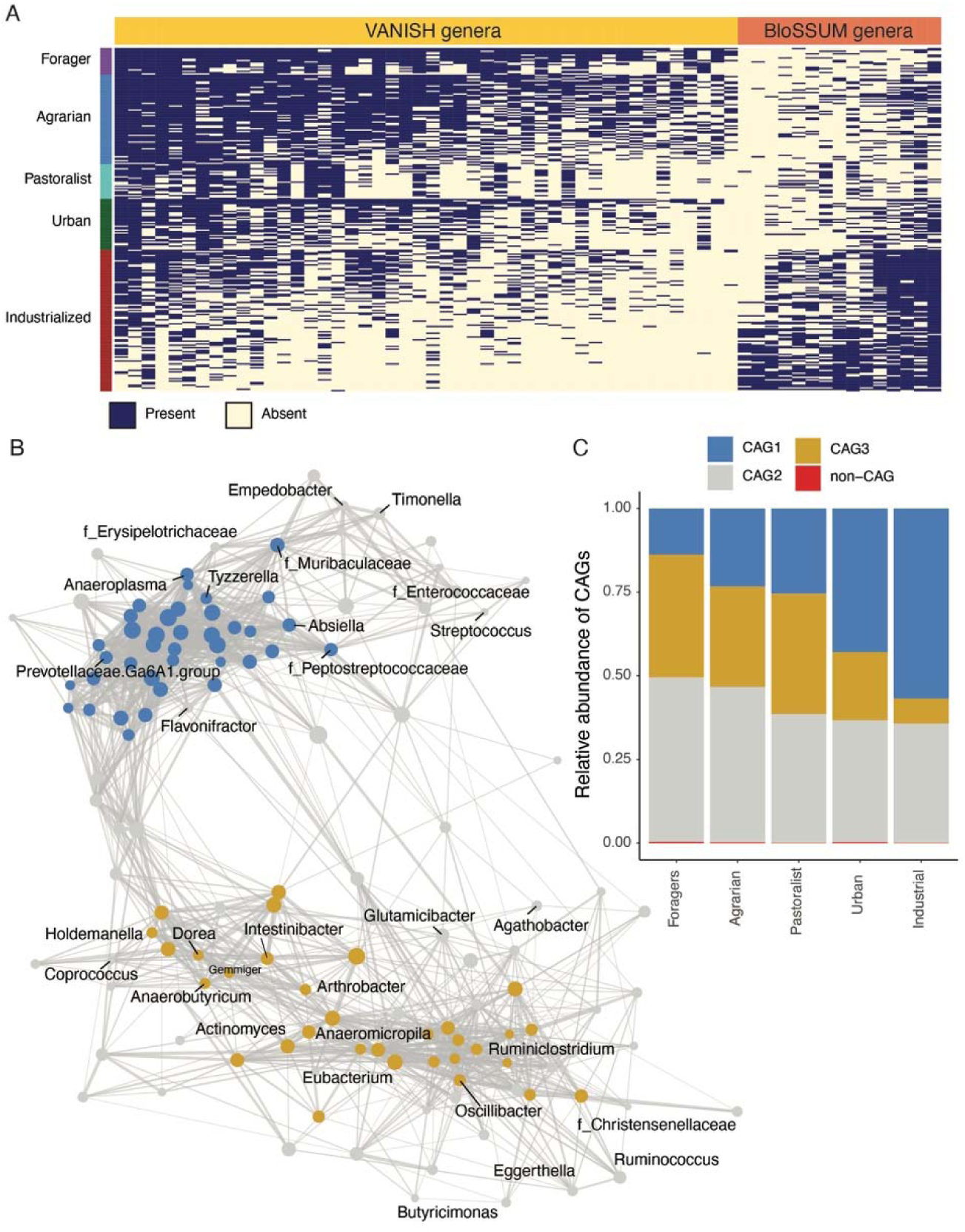
Prevalence-defined microbial guilds shift across the canine lifestyle gradient. **(A)** Presence-absence heatmap of VANISH-like and BloSSUM-like genera across dog samples ordered by lifestyle. VANISH-like and BloSSUM-like genera were identified by comparing prevalence between forager/agrarian and industrialized dogs using Fisher’s exact tests with FDR correction and requiring an absolute prevalence difference of at least 30%. Rows represent individual dogs and columns represent genera; navy indicates presence and pale yellow indicates absence. **(B)** Genus-level co-occurrence network inferred using FastSpar. Nodes represent genera and edges represent significant positive correlations. Node color indicates CAG membership, node size reflects mean genus abundance, and edge width reflects correlation strength. Key VANISH-like and BloSSUM-like genera are labeled. **(C)** Mean relative abundance of CAGs across lifestyle categories. Genera assigned to each CAG were summed within samples and converted to relative abundance across CAG and non-CAG taxa.

We then tested whether these prevalence-shifting genera were organized into coordinated co-abundance groups (CAGs). FastSpar [59] analyses resolved three CAGs, including two tightly connected modules and a third, more diffuse cluster (absolute rho > 0.2, FDR-adjusted P < 0.05; **Fig. S11**). VANISH-like and BloSSUM-like genera were non-randomly distributed across CAGs (P = 2.8 x 10^-7^, *Fisher’s exact test*; **Fig. 4B**). BloSSUM-like genera were confined to CAG1 and CAG2, whereas VANISH-like genera were restricted to CAG2 and CAG3. Consistent with this partitioning, CAG-level abundance varied directionally across lifestyles (**Fig. 4C**). CAG1, which was enriched for BloSSUM-like genera, increased toward industrialized dogs, whereas CAG3, composed primarily of VANISH-like genera, declined across the same gradient. Pastoralist dogs again showed a distinct non-industrialized profile, consistent with their departure from a simple forager-to-industrialized continuum. Together, these results indicate that industrialization in dogs is associated with coordinated turnover of prevalence-defined microbial guilds rather than isolated taxonomic shifts.

### Predicted functional potential mirrors the canine lifestyle gradient

We next tested whether lifestyle-associated taxonomic restructuring was accompanied by shifts in predicted functional potential inferred from 16S rRNA gene profiles using PICRUSt2 [60]. Because these profiles are predicted from marker-gene data rather than directly measured by shotgun metagenomics, we interpreted them as inferred functional potential.

Using a multivariable model including lifestyle, climate category, and age group, we identified predicted enzyme functions that followed the same monotonic forager-agrarian-urban-industrialized gradient used for genus-level analyses. This analysis identified 341 directional predicted enzyme functions (FDR-adjusted q < 0.05 and absolute MaAsLin2 coefficient > 1; **Table S8**). Most were enriched in forager dogs and declined toward industrialized dogs (307 functions), whereas a smaller set showed the reverse pattern (34 functions). These predicted functions spanned multiple enzyme classes, especially oxidoreductases, transferases, and hydrolases (**Fig. S12A**).

Predicted MetaCyc pathways showed a similar asymmetry. We identified 85 directional pathways enriched in forager dogs and declining toward industrialized dogs, whereas no pathways showed the reverse monotonic pattern under the same criteria (FDR-adjusted q < 0.05 and an absolute MaAsLin2 coefficient > 1; **Table S9**). These pathways spanned multiple MetaCyc superclass categories, including carbohydrate, lipid, and amino acid degradation and cofactor/vitamin biosynthesis (**Fig. S13**). Climate-associated predicted pathways were also detected, whereas age-associated functional gradients were comparatively modest (**Fig. S12B** and **Table S10**), consistent with the weaker age effects observed in taxonomic analyses. Overall, these results suggest that lifestyle-associated taxonomic restructuring is accompanied by broad shifts in predicted microbial functional potential. Because these profiles were inferred from 16S rRNA gene data, they should be interpreted as hypothesis-generating rather than as direct measurements of metagenomic function or metabolic activity.

### Shared lifestyle transitions produce host-specific microbiome responses in dogs and humans

We next asked whether lifestyle-associated microbiome restructuring in dogs paralleled patterns observed in humans sampled across similar lifestyle categories. We compared dogs and humans from matched forager, agrarian, and industrialized contexts using the human dataset from Jha et al. 2018 [5]. The human dataset included 44 individuals: 14 foragers, 20 agrarians, and 10 industrialized individuals. The matched dog dataset included 155 dogs: 20 foragers, 67 agrarians, and 68 industrialized dogs. Forager dogs and humans were sampled from the same village, strengthening the ecological comparability of the cross-host comparison.

Because dog and human samples were generated using different 16S rRNA amplicon regions, cross-host analyses were restricted to genus-level profiles. Separate Bray-Curtis PCoA analyses showed lifestyle-associated structure within each host: in both dogs and humans, industrialized samples separated from forager and agrarian samples along the leading ordination axis, whereas forager and agrarian samples were more similar to each other (**Fig. S14**). Thus, similar lifestyle categories were associated with broad community restructuring in both host species.

However, combined ordination of the 98 shared genera showed that host species remained the dominant axis of variation. PCA of CLR-transformed genus-level profiles separated dogs and humans along PC1, while lifestyle-associated differences occurred within host-specific compositional clusters (**Fig. 5A**). This pattern was robust to repeated stratified subsampling of dogs to match the number of human samples within each lifestyle category, indicating that host-specific separation was not driven by unequal sample size across hosts or lifestyles (**Fig. S15**). Thus, shared lifestyle transitions did not produce strong genus-level taxonomic convergence in overall community composition.

**Figure 5.**
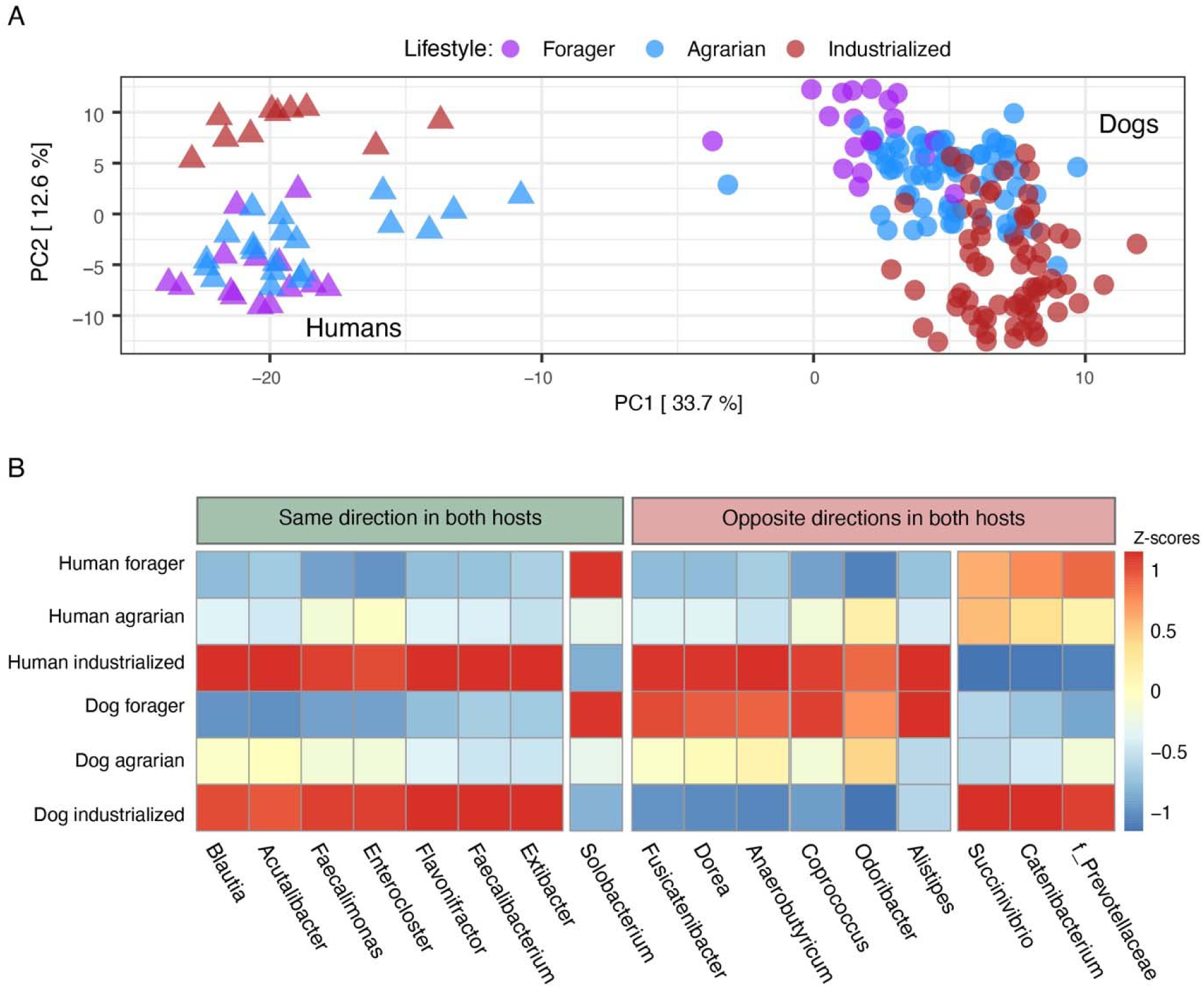
Shared lifestyle transitions produce host-specific microbiome responses in dogs and humans. **(A)** Principal component analysis (PCA) of centered log-ratio (CLR)-transformed genus-level abundance profiles after restricting the datasets to the 98 genera detected in both dogs and humans. Points represent individual samples, colored by lifestyle category and shaped by host species. Host species is the dominant axis of variation, while lifestyle-associated structure is evident within each host. **(B)** Heatmap of the 17 genera that showed significant directional lifestyle-associated responses in both dogs and humans, identified independently in each host. Columns show mean relative abundance across lifestyle groups within each host, scaled by row. Genera are grouped according to whether their lifestyle-associated responses occurred in the same direction or in opposite directions in dogs and humans.

We then tested whether the same genera responded to lifestyle transitions in both hosts. Among the 98 shared genera, 17 showed significant directional lifestyle-associated responses in both dogs and humans (FDR-adjusted q < 0.05 and absolute MaAsLin2 coefficient > 1; **Table S11**). Of these, only 8 changed in the same direction, whereas 9 changed in opposite directions (**Fig. 5B**). Same-direction overlap was not greater than expected under the shared genus background (P > 0.05, one-sided Fisher’s exact tests). These results indicate that lifestyle-associated restructuring occurred in both dogs and humans, but the genus-level taxa underlying these shifts were largely host-specific.

Together, the cross-host analyses show that industrialization and associated lifestyle transitions can impose parallel community-level pressures across human-associated mammals, but these pressures are expressed through host-specific microbial lineages rather than broad genus-level convergence.

## Discussion

Domestic dogs are a long-term human-associated mammal, but most canine gut microbiome studies have focused on industrialized companion animals [24, 25, 41, 61, 62]. This has limited our understanding of whether the microbiome of household pets represents a general canine baseline or one endpoint of a broader ecological spectrum. By analyzing dogs across forager, agrarian, pastoralist, urban, and industrialized contexts, we show that lifestyle is the strongest measured correlate of canine gut microbiome composition. Industrialized dogs were compositionally distinct from non-industrialized dogs and showed directional genus-level turnover, coordinated restructuring of VANISH- and BloSSUM-like microbial guilds, and shifts in inferred functional potential. At the same time, non-industrialized dogs were not microbiologically uniform: pastoralist dogs often departed from a simple forager-to-industrialized continuum, and climate contributed additional zone-specific variation. Restricted analyses further showed that lifestyle-associated structure persisted among mature, non-shelter dogs from temperate climate regions, and classifier-based projections placed Thai urban community dogs near the industrialized end of the microbiome gradient while assigning pastoralist dogs primarily to non-industrialized classes. These findings expand the ecological baseline for the domestic dog gut microbiome and identify industrialization as a major axis of microbiome restructuring in a long-term human-associated mammal.

A central finding is that lifestyle-associated variation in dogs was reflected in community composition rather than overall within-sample diversity. Alpha diversity showed no significant associations with measured host and ecological variables after multiple-testing correction, whereas beta diversity and genus-level composition differed strongly by lifestyle. Thus, industrialization-associated microbiome change in dogs should not be interpreted simply as loss of diversity. Instead, our results indicate that lifestyle transitions reorganize community structure by altering relative abundance and prevalence of bacterial lineages. This pattern is consistent with human studies showing that lifestyle-associated compositional differences can occur without uniform reductions in alpha diversity [5, 63].

The genus-level shifts across the canine lifestyle gradient suggest that industrialization is associated with both depletion and enrichment of distinct gut bacterial groups. Non-industrialized dogs showed higher abundance or prevalence of several anaerobic lineages previously associated with carbohydrate fermentation or short-chain fatty acid production, whereas industrialized dogs were enriched for other gut-associated genera, including taxa within Prevotellaceae, Erysipelotrichaceae, Lachnospiraceae, and Peptostreptococcaceae. These patterns likely reflect integrated differences in diet, environmental exposure, human management, and microbial transmission [64, 65]. However, because genus-level 16S data cannot resolve strain-level variation or functional heterogeneity within genera, these taxa should be interpreted as markers of lifestyle-associated community restructuring rather than direct evidence of specific metabolic roles.

Applying the VANISH/BloSSUM and co-occurrence analyses to dogs provided an additional link to the human industrialization literature [4, 58]. We observed analogous prevalence shifts in dogs: many VANISH-like genera were more prevalent in forager and agrarian dogs, whereas a smaller set of BloSSUM-like genera was more prevalent in industrialized dogs. These prevalence-shifting taxa were non-randomly distributed across co-abundance groups, and CAG-level abundance changed directionally across lifestyles. This suggests that industrialization-associated change in dogs involves coordinated microbial guilds rather than isolated taxonomic shifts. This is conceptually parallel to human industrialization patterns [5, 66]. However, the VANISH-like and BloSSUM-like categories describe prevalence shifts within the canine dataset and should not be assumed to represent the same bacterial lineages or ecological functions across host species. For example, several VANISH genera in humans such as *Segatella* and *Catenibacterium* that are often associated with carbohydrate- or fiber-rich diets in humans [67, 68], were enriched toward industrialized dogs. This may reflect host-specific nutritional ecology, commercial diet formulated with higher fiber ratio, or species-level heterogeneity within genus-level assignments.

The pastoralist dogs highlight why non-industrialized canine microbiomes should not be treated as a single “traditional” state. Pastoralist dogs shared free-ranging behavior and high environmental exposure with other non-industrialized dogs, but they occupied high-altitude settings and had distinct dietary and ecological contexts, including greater access to dairy and animal-derived foods [69]. Their separation along the secondary ordination axis suggests that local subsistence ecology can shape gut microbiomes in ways not captured by a simple industrialized versus non-industrialized contrast. Climate also contributed additional genus-level structure, especially through zone-specific taxa enriched in tropical environments. Because altitude and pastoralism were inseparable in this cohort, we interpret altitude as part of the integrated pastoralist ecological context rather than as an independent driver. The restricted classifier projection supported this interpretation: when forced into available forager, agrarian, urban, or industrialized labels, pastoralist dogs were assigned predominantly to the agrarian class and never to the industrialized class. Thus, pastoralist dogs appear to carry non-industrialized microbiome profiles while representing a distinct subsistence ecology rather than a simple intermediate stage. Together, these findings support treating lifestyle as an integrated ecological state that includes diet, human management, environmental exposure, climate, geography, and local subsistence ecology.

Predicted functional profiles broadly followed the lifestyle gradient, with many inferred enzyme functions and pathways enriched in forager dogs and declining toward industrialized dogs. This pattern is consistent with the observed taxonomic restructuring and suggests that lifestyle-associated community shifts may be accompanied by changes in metabolic potential. However, these profiles were inferred from 16S rRNA gene data rather than measured by shotgun metagenomics or metabolomics and therefore should be interpreted as supportive evidence of predicted functional differences, not as direct evidence of metabolic activity. Future metagenomic and metabolomic studies will be needed to test whether the taxonomic patterns observed here correspond to functional differences in carbohydrate degradation, lipid metabolism, amino acid metabolism, and microbial metabolite production.

The dog-human comparison places these findings in a broader comparative context. Prior studies show that shared ecology can increase microbiome similarity across host species, but that convergence is usually partial and context-dependent. Geographic coexistence, captivity, urbanization, and strong dietary specialization can all increase cross-host similarity [32, 34-40, 70], yet host-specific structure generally persists. Dogs therefore provide a particularly stringent natural test of whether shared ecology can overcome host specificity because they have coexisted with humans over long timescales and across shared lifestyle transitions. Our results show that dogs and humans both show lifestyle-associated microbiome restructuring across forager, agrarian, and industrialized categories. However, combined analyses of shared genus-level taxa showed that host species remained the dominant axis of variation, and lifestyle-associated shifts occurred within host-specific compositional space rather than producing strong cross-host convergence. The genus-level taxa responding to lifestyle transitions were also largely host-specific, and same-direction overlap was not greater than expected under the shared-genus background. This pattern remained robust after stratified subsampling of dogs to match the number of human samples within each lifestyle category. Thus, similar lifestyle transitions can impose parallel community-level pressures across host species, but the microbial lineages through which those pressures are expressed remain strongly shaped by host biology.

This result clarifies the value of dogs as a comparative microbiome system. Dogs are not simple microbiome proxies for humans, despite their long history of cohabitation, shared environments, and consumption of human-associated foods. Instead, they provide a natural system for separating shared ecological pressure from shared microbial outcome. The absence of strong genus-level dog-human taxonomic convergence indicates that host physiology, immune filtering, gastrointestinal traits, diet processing, transmission history, and evolutionary background can constrain how industrialization-associated pressures are expressed. In this sense, dogs reveal both the reach and the limits of shared human ecology: industrialization can restructure gut microbiomes across human-associated mammals, but it does not erase host-specific community assembly.

Several limitations should be considered. First, this was an observational study across natural ecological gradients, where lifestyle, geography, climate, altitude, and human management are inherently coupled. We therefore interpret lifestyle as an integrated ecological exposure rather than as a single causal variable. Restricted analyses showed that lifestyle-associated structure persisted after limiting the dataset to mature, non-shelter dogs from temperate climate regions and excluding Thai and UAE dogs, but these analyses cannot fully disentangle all components of lifestyle or exclude unmeasured covariates. Second, 16S rRNA gene sequencing limits taxonomic resolution and does not capture strain-level variation, gene content, or directly measured functional activity. Third, dietary information was categorical and incomplete for some populations, particularly dogs from Thailand. Although Thai dogs projected near the industrialized end of the microbiome gradient, they should be interpreted as urban community dogs with potential processed-food exposure rather than industrialized household pets. Fourth, dog-human comparisons were limited by differences in host biology, sample size, and 16S rRNA gene amplicon region, requiring genus-level harmonization. Stratified subsampling supported the robustness of the host-specificity result, but paired dog-human sampling, shotgun metagenomics, and metabolomics would provide a stronger test of taxonomic, strain-level, and functional sharing.

## Conclusions

Our findings expand the ecological baseline for the domestic dog gut microbiome beyond industrialized companion animals and show that industrialization is associated with robust, directional restructuring of canine gut microbial communities. The dog-human comparison further indicates that shared lifestyle transitions can produce parallel microbiome restructuring without strong genus-level taxonomic convergence, highlighting domestic dogs as a useful comparative system for studying how human-associated ecological change shapes mammalian gut microbiomes while preserving host-specific community structure.

## Methods

### Ethics approval and consent to participate

All animal procedures were approved by the New York University Institutional Animal Care and Use Committee (IACUC Protocol #20-0005) and the Nepal Veterinary Council (Reg #535). Fecal samples were collected non-invasively from freshly voided feces or during routine handling with permission from owners, caretakers, or responsible community members where applicable. No animals were subjected to experimental procedures or interventions beyond routine handling for sample collection. Sample collection, transport, storage, and data sharing complied with applicable animal research, biosafety, and material transfer regulations.

### Optimization of fecal sample preservation

To select a field-compatible preservation method, we compared three commercial fecal preservation systems with flash-frozen reference samples using fecal samples from seven pet dogs at New York University Abu Dhabi. Microbial profiles differed between flash-frozen and preserved samples, whereas differences among preservation systems were modest (**Fig. S16**). We selected the Huechenyan fecal collection kit for field sampling because it performed comparably to the other preservation systems and was substantially less expensive.

### Sample collection

Dog fecal samples were collected across Nepal, Thailand, the UAE, and the United States to capture variation in lifestyle, climate, geography, altitude, and human association. In Nepal, Huechenyan fecal collection kits were used to collect fecal samples from 151 dogs across nine communities representing four non-industrialized lifestyle categories. Twenty forager dogs were sampled from Chepang communities in Korak at approximately 1,400 m elevation; 67 agrarian dogs were sampled in Chobhar, Parsa, and Nuwakot at 100-1,500 m elevation; 26 pastoralist dogs were sampled from high-altitude regions of Rasuwa, Tsarang, and Lo-Manthang at 2,000-3840 m elevation; and 38 urban dogs were sampled in Chitwan and Banepa at 100-1,500 m elevation. Most Nepali samples were obtained from free-ranging community dogs, except for 15 dogs sampled at the Chobhar rescue shelter. None of the Nepali dogs were reported to consume commercial pet food; instead, they consumed household and scavenged foods.

Outside Nepal, dogs were sampled in Chiang Mai, Thailand (n = 27); Abu Dhabi, UAE (n = 13); and the United States (n = 70), including Louisiana (n = 5), New York (n = 11), Pennsylvania (n = 15), and North Carolina (n = 39). Dogs from Chiang Mai were sampled in urban settings and included free-ranging community dogs brought to a rescue center and sampled on the day of rescue. Their individual diets prior to rescue were not known with confidence, but community dogs in Chiang Mai may forage within an urbanizing food environment where traditional diets coexist with greater access to commercial and processed foods [47]. Thai dogs were therefore treated as urban community dogs with potential exposure to processed human food waste, rather than industrialized household pets. Nine dogs from North Carolina were sampled at an animal shelter, whereas the remaining UAE and US dogs were household pets consuming commercial pet food.

Dogs from Chiang Mai, Abu Dhabi, Louisiana, New York, and North Carolina were sampled using the Huechenyan fecal collection kits. Fecal samples from Pennsylvania pet dogs were collected on ice, transported to the laboratory, and frozen. All international shipments were transported on dry ice, and samples were stored at -80 °C until DNA extraction. Samples were processed and sequenced together using a standardized protocol to minimize technical variation.

Dogs were assigned to tropical, arid-desert, temperate, and temperate-cold climate categories based on sampling location using Koppen-Geiger climate classification [71]. ET and Df zones were grouped as temperate-cold because both corresponded to cooler, non-arid environments in this dataset. Dogs were categorized as young (<0.5 years), mature (0.5-7 years), and older (>7 years), following veterinary life-stage guidelines [72]. Metadata including climate category, altitude, age group, and recent antibiotic exposure, when available, are provided in **Table S1**.

### DNA extraction and 16S rRNA gene sequencing

Samples were mechanically lysed using a Bead Ruptor 96 bead mill (Omni International, USA), followed by DNA extraction with the ZymoBIOMICS PowerSoil DNA Extraction Kit (D4303). Extractions were performed in individual tubes, and negative extraction controls were included after every eight samples.

A total of 351 samples, including 309 dog fecal samples and 42 negative extraction controls, yielded sufficient DNA for sequencing. Extracted DNA was shipped on dry ice to Novogene Singapore for library preparation and sequencing. The V3-V4 region of the bacterial 16S rRNA gene was amplified using primers 341F/806R with barcode-tagged forward primers [73]. Twenty-nine negative controls and 20 fecal samples failed to amplify. PCR products from the successfully amplified samples, comprising 289 fecal samples and 13 controls, were purified using AMPure XP beads, quantified using the Qubit dsDNA BR Assay kit, pooled, and sequenced on an Illumina platform using paired-end 2 × 300 bp chemistry.

### Amplicon sequence processing

Sequencing read quality was assessed using FastQC, and amplicon sequence variants (ASVs) were inferred using *DADA2* [74]. Reads were filtered and trimmed using maxN = 0, maxEE = 2, truncQ = 2; error rates were learned from the data, ASVs were inferred using pooled processing, paired-end reads were merged, and chimeras were removed. Taxonomy was assigned using the RDP classifier [75] using the SILVA v138 training set [76]. ASVs classified as Eukaryota, chloroplast, mitochondria or unclassified at the phylum level were removed and ASVs represented by more than 10 reads in more than five samples were retained. ASVs were aligned using *DECIPHER* [77], and a phylogenetic tree was constructed using *phangorn* [78].

Downstream analyses were conducted in R v4.2.0 using *phyloseq* [79]. Among the 302 sequenced samples, three duplicate fecal samples were removed. Thirteen controls passed sequencing quality filters and were retained for contamination assessment. To identify contamination-like or low-biomass samples, we performed PCoA and partitioning around medoids (PAM) clustering including all negative controls; 15 fecal samples clustered with controls **(Fig. S17)**. Together with the 13 retained controls, these samples were excluded from downstream analyses. Ten additional fecal samples with fewer than 5,000 reads were removed. After filtering, 261 fecal samples remained, comprising 8,763,965 reads and 1,472 ASVs assigned to 245 bacterial genera. Four dogs with reported recent antibiotic exposure were excluded from downstream microbiome analyses, yielding 257 dogs in the final dataset.

### Beta diversity analyses

Beta diversity was assessed using weighted UniFrac and Bray-Curtis distances calculated from genus-level abundance data after log(x + 1) transformation as described previously [80]. Weighted UniFrac distances were calculated in *phyloseq* using the phylogenetic tree with normalization enabled. Principal coordinate analysis (PCoA) was performed using *phyloseq* and visualized with *ggplot2* [81]

Associations between gut microbiome composition and host or ecological metadata were tested using PERMANOVA with *adonis2* in the *vegan* package [82] and 999 permutations. Multivariable models included lifestyle, climate category, altitude category, shelter status, and age group with marginal effects estimated using *by = “margin”.* EnvFit analyses were performed using the first three ordination axes with 999 permutations to assess alignment of metadata variables with major compositional gradients.

To assess robustness to major measured confounders, beta-diversity analyses were repeated in a restricted subset of mature, non-shelter dogs sampled from temperate climate regions, excluding Thai and UAE dogs. This restricted subset was tested using the same weighted UniFrac and Bray-Curtis distance metrics. PERMANOVA models included lifestyle as the predictor variable, and EnvFit analyses assessed alignment of lifestyle with the first three ordination axes.

### Random forest classifiers

Random forest classifiers were trained on log(x + 1)-transformed genus-level abundance profiles using the *tidymodels* framework [83] with the *ranger* engine to predict lifestyle category. Models were generated for the full 257-dog dataset and for the restricted mature, non-shelter, temperate-climate subset. The full-dataset model was used only as a supplemental performance comparison. The restricted classifier was then used for projection of excluded dogs.

Samples were partitioned into training and held-out test sets using an 80/20 partition. Within the training set, five-fold cross-validation and grid search were used to tune *mtry* and *min_n*; models were trained with 500 trees. Final models were evaluated on held-out test sets using accuracy, Cohen’s kappa, and multiclass ROC AUC calculated from one-vs-all class probabilities with the *yardstick* package [84].

For the restricted sensitivity and projection analysis, the classifier was trained using only mature, non-shelter dogs sampled from temperate climate regions, excluding Thai and UAE dogs. This restricted training set included 107 dogs spanning forager, agrarian, urban, and industrialized lifestyles. Pastoralist dogs were not included as a training class because they represented a distinct high-altitude ecological context rather than a stage along the forager-to-industrialized gradient. The classifier was trained on log(x + 1)-transformed genus-level abundance profiles using the same modeling framework described above. Model performance was evaluated using an 80/20 train-test split. The trained classifier was then used to project dogs excluded from the restricted training set, including Thai, UAE, additional US, excluded Nepali, and pastoralist dogs. Projection results were summarized by country and observed lifestyle category. Balloon plots show row-wise proportions assigned to each predicted lifestyle class, with sample counts shown inside points.

### Alpha diversity analyses

Alpha diversity was quantified as observed ASV richness, Shannon diversity, and Faith’s phylogenetic diversity. To assess sensitivity to sequencing depth, ASV tables were rarefied from 1,000 to 6,000 reads in 1,000-read increments, with 10 iterations per depth. Rarefaction curves indicated saturation at 6,000 reads; therefore, downstream alpha diversity analyses used estimates calculated at a rarefaction depth of 6,000 reads. Associations between alpha diversity and metadata were tested using generalized linear models with a Gamma distribution and log link, including lifestyle, climate category, shelter status, and age group as predictors. Altitude category was not included because it was collinear with climate category in this dataset.

### Differential abundance

Genus-level differential abundance was tested using MaAsLin2 [48]. Two models were used. First, a lifestyle-only model was fit with industrialized dogs as the reference group to identify genera associated with the primary lifestyle gradient. Second, a multivariable model including lifestyle, climate category, and age group as fixed effects was fit to assess whether lifestyle-associated genera remained detectable after accounting for major ecological and host covariates. In this model, industrialized, tropical, and young dogs were used as reference populations for lifestyle, climate, and age, respectively.

Both models used linear models with cumulative sum scaling normalization, log transformation, and a minimum prevalence threshold of 5%. Genera were considered significant at FDR-adjusted q value < 0.05. Large-effect associations were defined as FDR-adjusted q < 0.05 with absolute MaAsLin2 coefficient > 1, corresponding approximately to a greater than twofold difference in normalized abundance on the log-transformed scale.

To identify directional lifestyle-associated taxa, we applied a monotonic-gradient criterion across the forager-agrarian-urban-industrialized axis. Genera were retained if coefficients followed either a decreasing pattern from forager to agrarian to urban to industrialized dogs or the reverse pattern, and if the forager-versus-industrialized endpoint contrast met the large-effect threshold. Pastoralist dogs were excluded from this monotonic-gradient because they represented a distinct high-altitude lifestyle rather than a simple intermediate stage along the forager-to-industrialized axis.

### Identification of VANISH- and BloSSUM-like genera

VANISH- and BloSSUM-like genera were identified using a prevalence-based comparison between non-industrialized and industrialized dogs. For this analysis, forager and agrarian dogs were grouped as the non-industrialized reference set and compared with industrialized dogs. Genus-level abundances were converted to presence/absence, and prevalence differences between groups were tested using Fisher’s exact test with Benjamini-Hochberg correction.

Genera were classified as VANISH-like if they were significantly more prevalent in non-industrialized dogs, and as BloSSUM-like if they were significantly more prevalent in industrialized dogs. To focus on robust prevalence shifts, genera were retained if they had FDR-adjusted q < 0.05 and an absolute prevalence difference ≥ 0.30.

### Co-abundance group analyses

Co-abundance groups (CAGs) were inferred from genus-level abundance profiles using FastSpar [59] and permutation-based p-values were estimated with 1,000 bootstrap. Pairwise associations were filtered to retain genus pairs with FDR-adjusted P < 0.05 and absolute correlation coefficient > 0.2. Genera involved in at least one significant association were clustered by hierarchical clustering of the filtered correlation matrix, yielding three major CAGs.

For visualization, positive genus-genus associations were plotted as a network plot using *igraph* [85]. To quantify CAG abundance across lifestyles, genera assigned to each CAG were summed within each sample and remaining genera were grouped as a non-CAG fraction. CAG abundances were converted to sample-wise relative abundances and summarized by lifestyle. Enrichment of VANISH- and BloSSUM-like genera across CAGs was tested using Fisher’s exact test, with P < 0.05 used as the significance threshold.

### Predicted functional profiling and pathway annotation

Predicted functional profiles were generated from 16S rRNA gene data using PICRUSt2 [60] and interpreted as inferred functional potential rather than directly measured metagenomic function. EC-numbered enzyme features and MetaCyc pathway abundance tables were analyzed separately, retaining features present in at least 10% of samples.

Associations between predicted functional profiles and host/ecological variables were tested using MaAsLin2 with total-sum scaling normalization, log transformation, and linear modeling. Models included lifestyle, climate category, and age group as fixed effects, with industrialized dogs specified as the lifestyle reference group, tropical as the climate reference group, and young dogs the age reference group. Significant associations were defined as FDR-adjusted q < 0.05, and large-effect associations were defined as FDR-adjusted q < 0.05 with absolute MaAsLin2 coefficient > 1.

Directional lifestyle-associated predicted functions were identified using the same monotonic-gradient criterion applied to genus-level taxa across the forager-agrarian-urban-industrialized axis. Features were retained if coefficients followed either a decreasing pattern from forager to industrialized dogs or the reverse pattern, and if the forager-versus-industrialized endpoint contrast met the large-effect threshold. Pastoralist dogs were excluded from this monotonic-gradient definition. EC features were assigned to broad enzyme classes using the first digit of the EC identifier, and MetaCyc pathways were annotated using MetaCyc pathway classes [86].

### Cross-species comparisons between dogs and humans

Dog data from this study and human data from Jha et al. 2018 [5] were restricted to lifestyle categories shared across both datasets: foragers and agrarians from Nepal and industrialized samples from the United States. The human dataset included 44 individuals: 14 foragers, 20 agrarians, and 10 industrialized participants. The matched dog dataset included 155 dogs: 20 foragers, 67 agrarians, and 68 industrialized dogs. Raw human FASTQ files were reprocessed using the same *DADA2* workflow and taxonomic assignment strategy applied to the dog dataset.

Because human samples were profiled using the V4 region of the 16S rRNA gene and dog samples using the V3-V4 region, cross-host analyses were performed at the genus level rather than at the ASV level. A total of 98 genera were detected in both datasets. Dog and human datasets were first analyzed separately using Bray-Curtis PCoA of genus-level abundance profiles to assess lifestyle-associated structure within each host. Genus-level differential abundance analyses were then performed separately in dogs and humans using MaAsLin2, with industrialized samples specified as the reference group. Directional lifestyle-associated genera were identified using the same monotonic-gradient criterion applied in the dog analysis.

For combined dog-human compositional analyses, datasets were restricted to the 98 shared genera. Zero replacement was performed using the count zero multiplicative method in *zCompositions*, followed by centered log-ratio transformation. Principal component analysis was then applied to the CLR-transformed genus-level matrix to visualize major axes of variation across hosts and lifestyles.

To test whether lifestyle-associated genera overlapped between host species, the 98 shared genera were used as the background. Same-direction overlap was assessed separately for genera enriched in forager-associated microbiomes and genera enriched in industrialized microbiomes using one-sided Fisher’s exact tests.

### Robustness of dog-human PCA patterns to matched subsampling

To assess whether cross-host PCA patterns were affected by unequal sample sizes, we repeated the dog-human comparison after stratified subsampling of dogs to match the number of human samples in each lifestyle category: 14 foragers, 20 agrarians, and 10 industrialized. In each of 100 iterations, dog samples were randomly selected without replacement within each lifestyle category, merged with the full human dataset using shared genera, and PCA was repeated using the same genus-level abundance transformation as in the primary analysis. For each iteration, we extracted sample-level PCA coordinates, variance explained, and host x lifestyle centroids. Robustness was evaluated from the stability of centroid positions across iterations.

## Supplementary Information

Supplementary Figures include Figure S1-S17. Supplementary Tables include Table S1-S11.

## Availability of data and material

Summary data necessary to reproduce the analyses presented in this manuscript are provided in the Supplementary Tables. Raw amplicon sequencing data have been deposited in the Sequence Read Archive under accession PRJNA1472931. A phyloseq object containing sample IDs, host-associated metadata, ASV-level count data, taxonomic annotations, and the phylogenetic tree is also included as Supplementary Data File. Additional information needed to reanalyze the data is available from the corresponding author upon reasonable request.

## Competing interests

The authors declare no competing interests.

## Funding

This work was supported by the Startup Grant (Grant No. ST318) and Faculty Research Grant from New York University Abu Dhabi to A.R.J. (Grant No. AD318).

## Authors’ contributions

A.R.J. conceived the study. Ar.G., D.B., Aa.G., P.Y., K.C.M.S., and K.A.H. performed field work and sample collection. Ar.G. and A.R.A. performed experiments. Ar.G., Ku.G., Ks.G., and A.R.J. performed data analyses. Ar.G., D.B., Ku.G., K.C.M.S., K.A.H., L.S.W., A.K.K., and A.R.J. interpreted the results. Ar.G., Ku.G., and A.R.J. generated the figures. D.S., K.A.H., L.S.W., A.K.K., A.R.J. supervised the project. Ar.G., Ku.G., and A.R.J. wrote the manuscript with input from all authors. All authors approved the final manuscript.

## Acknowledgments

We thank all community members in Nepal, Thailand, the UAE, and the United States who facilitated sample and data collection, including Jung Kunwar, Bimarsh Rana, and Yadhusudan Kunwar. We thank Novogene for sequencing services and the NYUAD High-Performance Computing Core for computational resources. We are grateful to members of Genetic Heritage Group and Sabitri Sciences for constructive discussions throughout this study. We also thank New York University Institutional Animal Care and Use Committee (IACUC) and the Nepal Veterinary Council for providing the necessary approvals and permits. We are grateful to Emily Davenport and Matt Olm for helpful discussions and critique of the manuscript.

## Supplementary Information

### Additional File 1. Supplementary Figures

**Figure S1.**
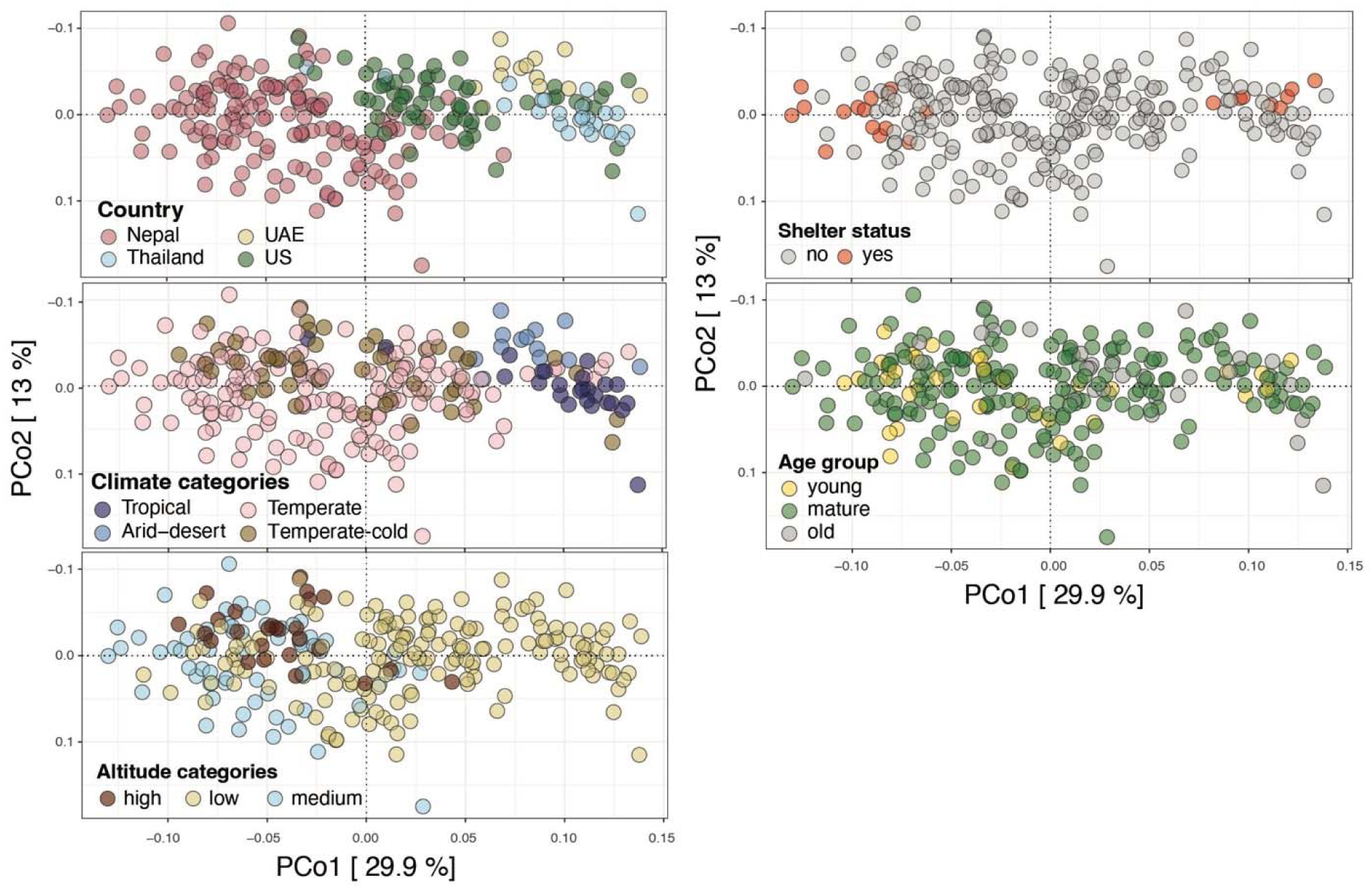
Ecological and host metadata projected onto weighted UniFrac PCoA space. Weighted UniFrac PCoA of genus-level microbiome profiles from 257 dogs, colored separately by country, climate category, altitude category, shelter status, and age group. Axes correspond to the same ordination shown in Fig. 2A. These projections show the partial coupling among lifestyle-associated ecological variables, including country, climate, and altitude, while shelter status and age show weaker structure.

**Figure S2.**
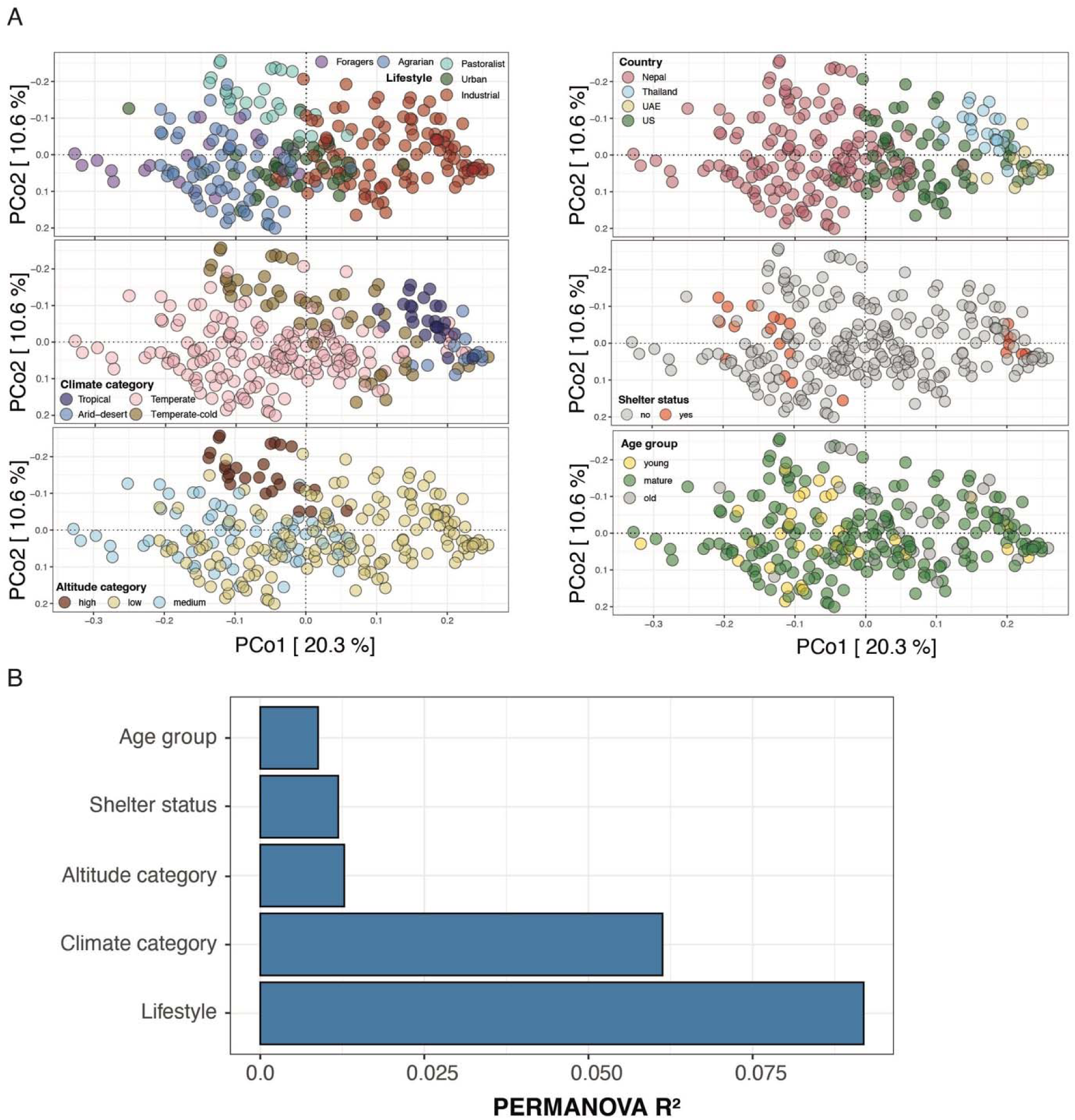
Bray-Curtis analyses support lifestyle-associated microbiome structure. **(A)** Bray-Curtis PCoA of genus-level microbiome profiles from 257 dogs, colored separately by lifestyle, country, climate category, altitude category, shelter status, and age group. Axes show the first two principal coordinate axes, with percent variation explained indicated in brackets. **(B)** Marginal PERMANOVA R² values from a multivariable model testing associations between Bray-Curtis composition and lifestyle, climate category, altitude category, shelter status, and age group. Results show that lifestyle explained the largest share of Bray-Curtis variation, followed by climate category, consistent with the weighted UniFrac analyses.

**Figure S3.**
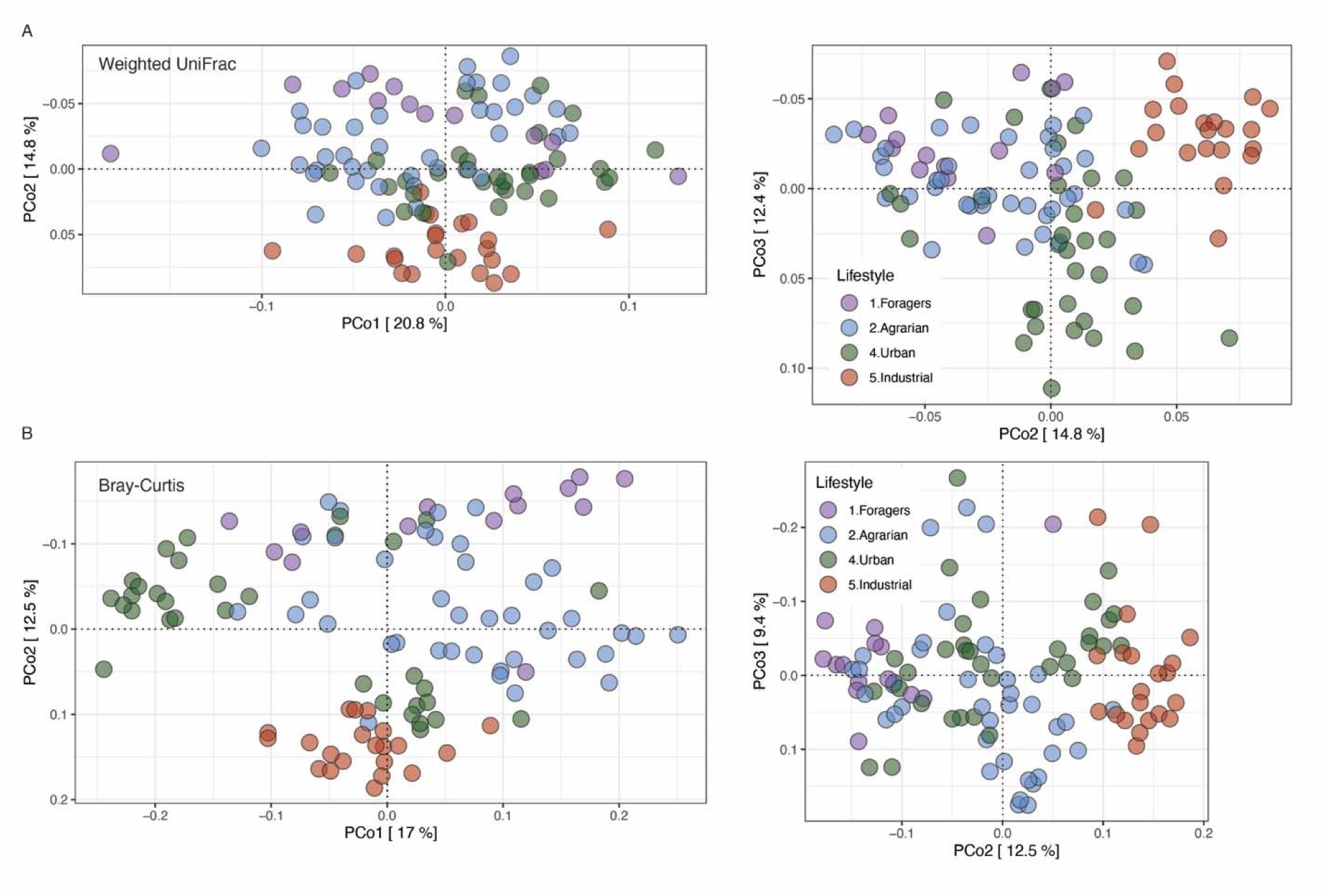
Lifestyle-associated microbiome structure persists in a restricted subset of mature, non-shelter, temperate-climate dogs. Weighted UniFrac and Bray-Curtis analyses were repeated in a restricted subset of 107 mature, non-shelter dogs sampled from temperate climate regions, excluding Thai and UAE dogs. This subset retained forager, agrarian, urban, and industrialized dogs. Lifestyle remained strongly associated with microbiome composition in both weighted UniFrac and Bray-Curtis analyses, explaining nearly 20% of variation in each distance metric (weighted UniFrac: R² = 0.197, P = 0.001; Bray-Curtis: R² = 0.196, P = 0.001; PERMANOVA; Table S2).

**Figure S4.**
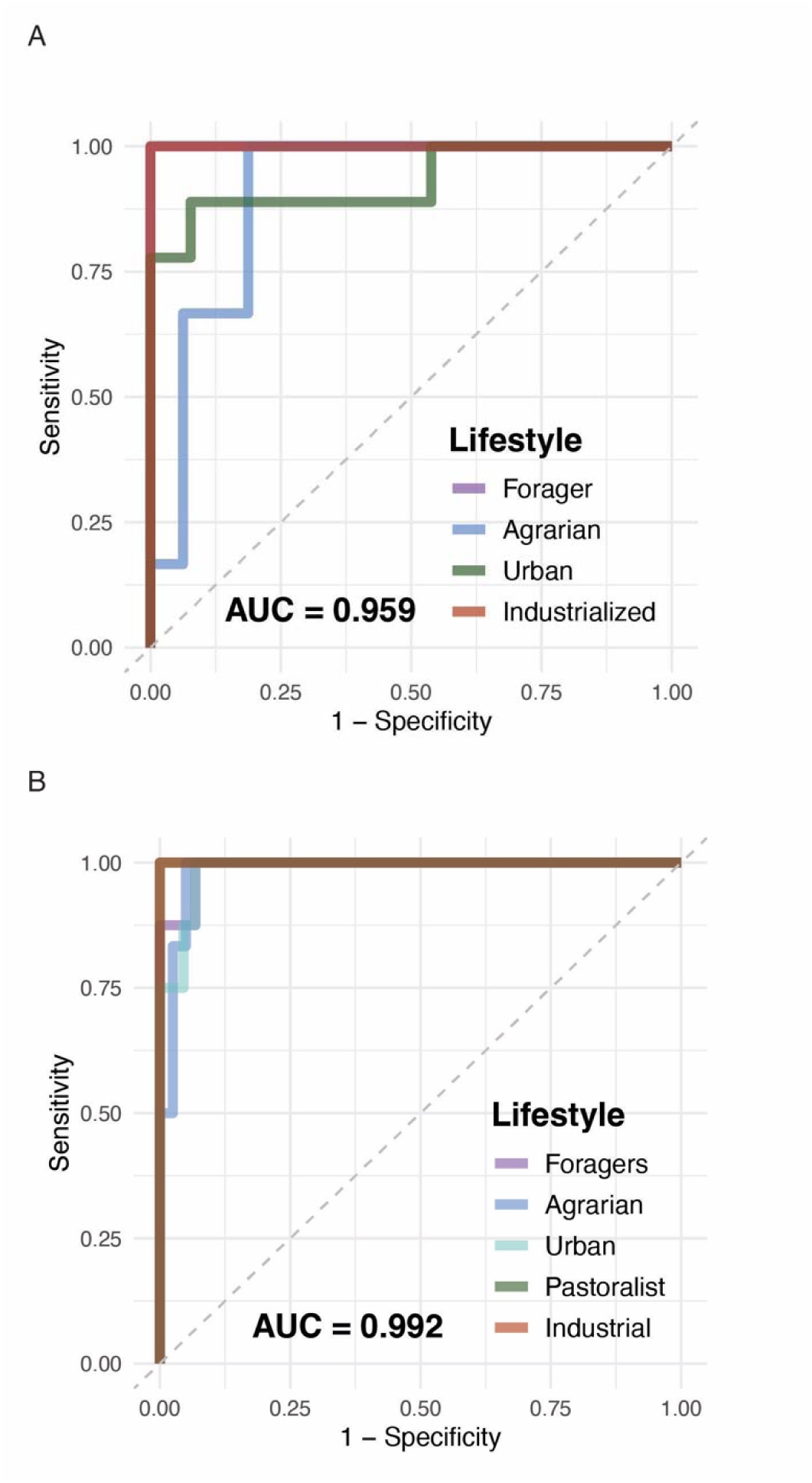
Random forest classification performance for restricted and full canine microbiome datasets. Multiclass receiver operating characteristic curves for random forest classifiers trained on log(x + 1)-transformed genus-level microbiome profiles. Curves show one-vs-all classification for each lifestyle class, and AUC values indicate macro-average AUC across classes. **(A)** Restricted classifier trained on 107 mature, non-shelter dogs sampled from temperate climate regions, excluding Thai and UAE dogs; pastoralist dogs were excluded from training because they represented a distinct high-altitude ecological context rather than a stage along the forager-to-industrialized gradient. **(B)** Classifier trained on the full dataset of 257 dogs after excluding individuals with recent antibiotic exposure.

**Figure S5.**
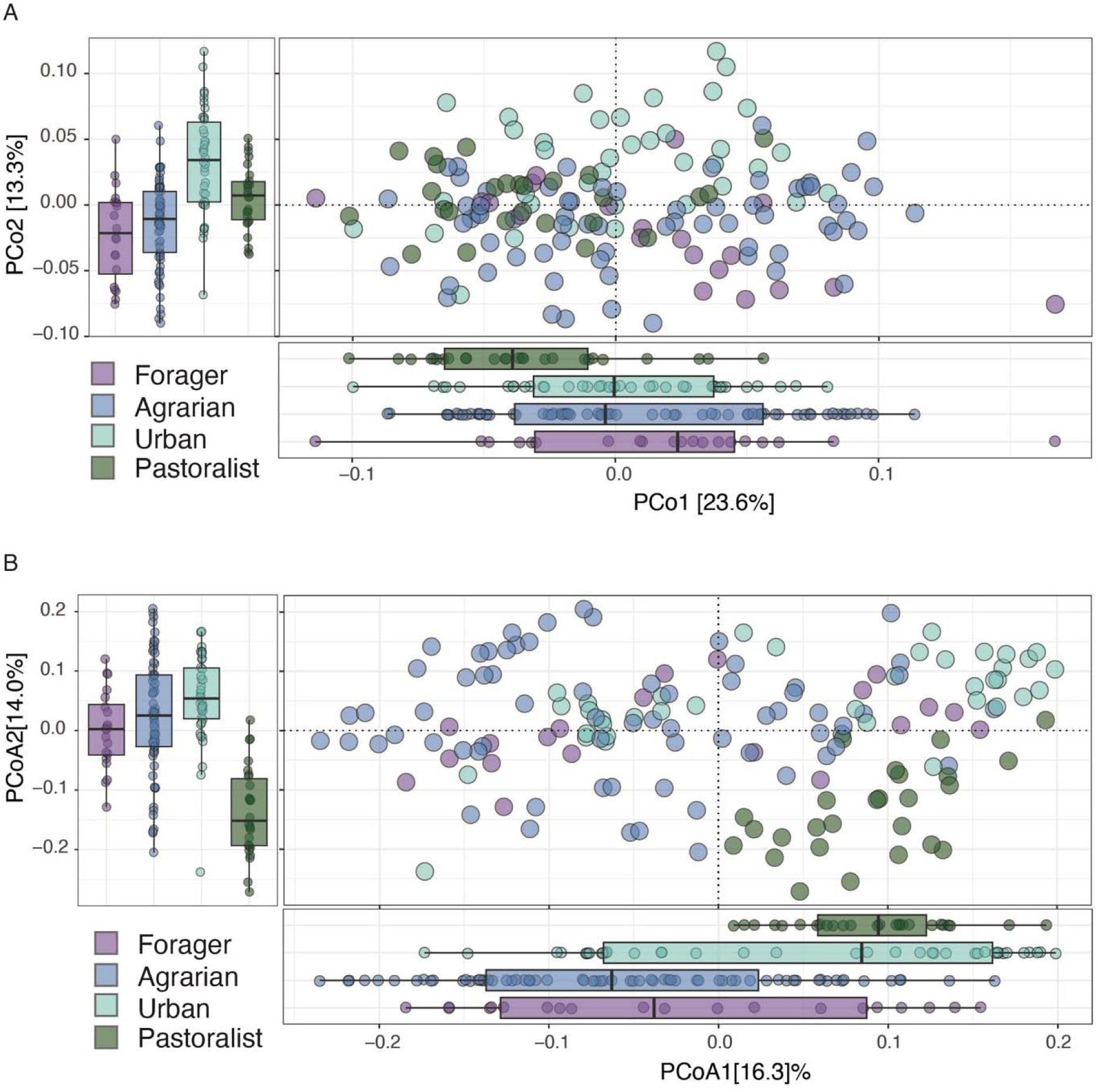
Lifestyle-associated microbiome structure within Nepal. **(A)** Weighted UniFrac PCoA of genus-level microbiome profiles from Nepali dogs only, colored by lifestyle category. Marginal boxplots show the distribution of samples along PCo1 and PCo2. **(B)** Bray-Curtis PCoA of the same Nepali subset, colored by lifestyle category. Lifestyle-associated structure remained evident among Nepali dogs alone, with forager dogs at one end of the ordination, pastoralist dogs at the other, and agrarian and urban dogs occupying intermediate positions.

**Figure S6.**
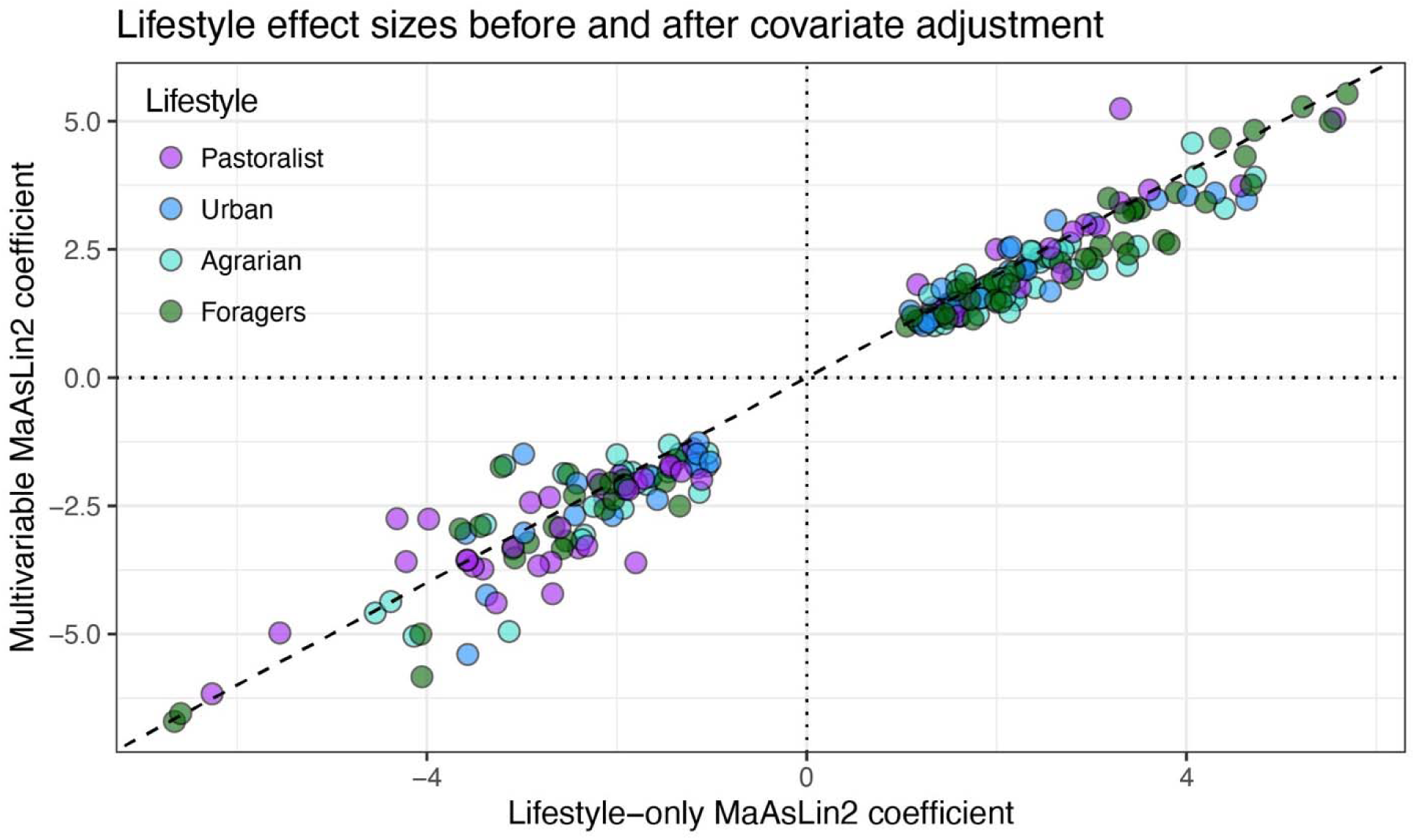
Lifestyle-associated genus-level effect sizes are preserved after covariate adjustment. Scatterplot comparing MaAsLin2 coefficients from the lifestyle-only model (x-axis) and the multivariable model including lifestyle, climate category, and age group (y-axis) across shared genus-lifestyle contrasts. Points are colored by lifestyle contrast relative to industrialized dogs. The dashed line indicates the 1:1 relationship, and dotted lines mark zero effect. Effect directions were fully concordant between models, and coefficients were strongly correlated (r = 0.98), indicating that the magnitude and direction of lifestyle-associated genus-level shifts were largely preserved after adjustment for covariates.

**Figure S7.**
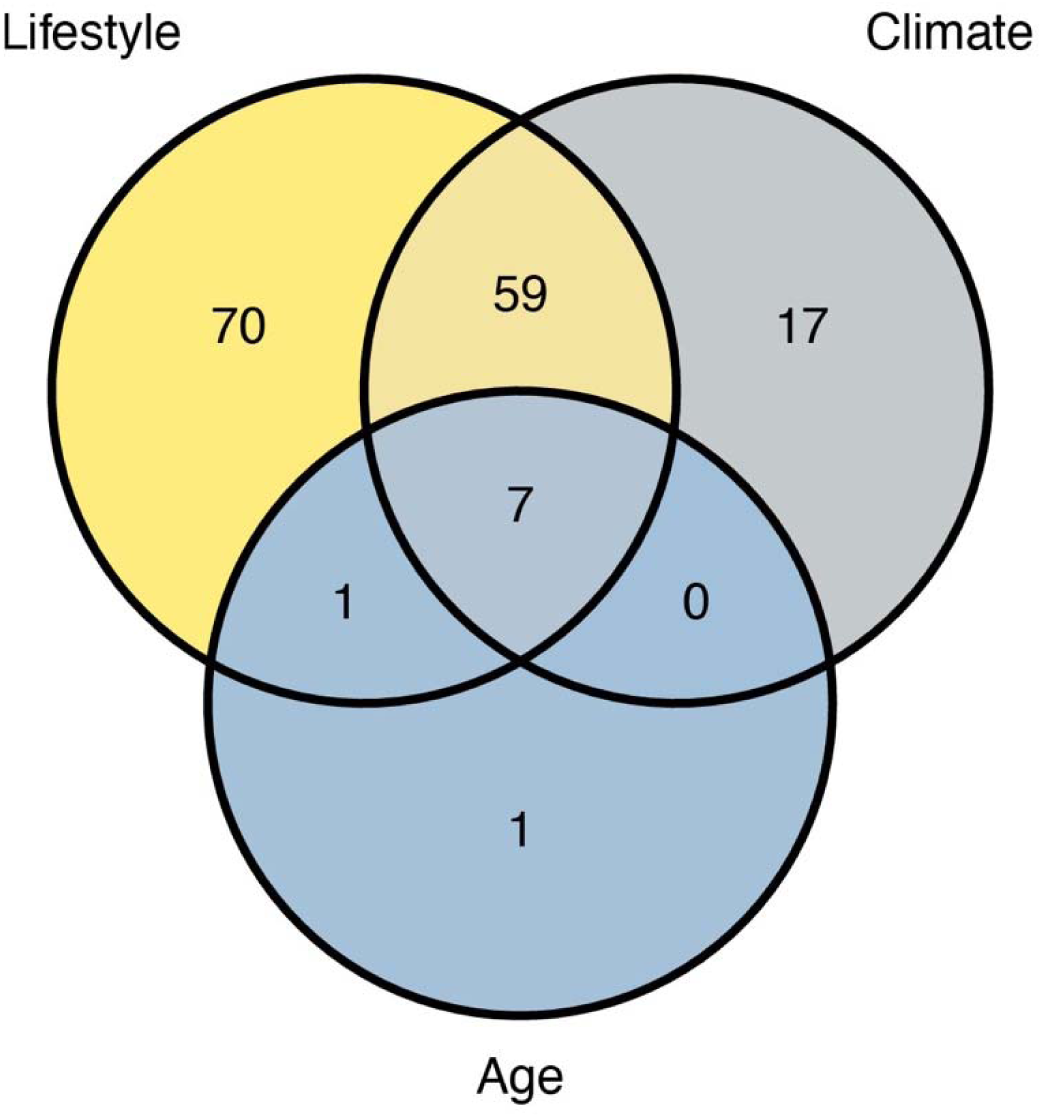
Overlap among lifestyle-, climate-, and age-associated genera. Venn diagram showing overlap among genera with significant large-effect associations with lifestyle, climate category, or age group in the multivariable MaAsLin2 model. Genera were included if they showed FDR-adjusted q < 0.05 and absolute MaAsLin2 coefficient > 1 for at least one contrast within a metadata category. Lifestyle and climate associations partially overlapped, but many lifestyle-associated genera were not associated with climate category, indicating that lifestyle captured substantial genus-level variation beyond broad climatic differences.

**Figure S8.**
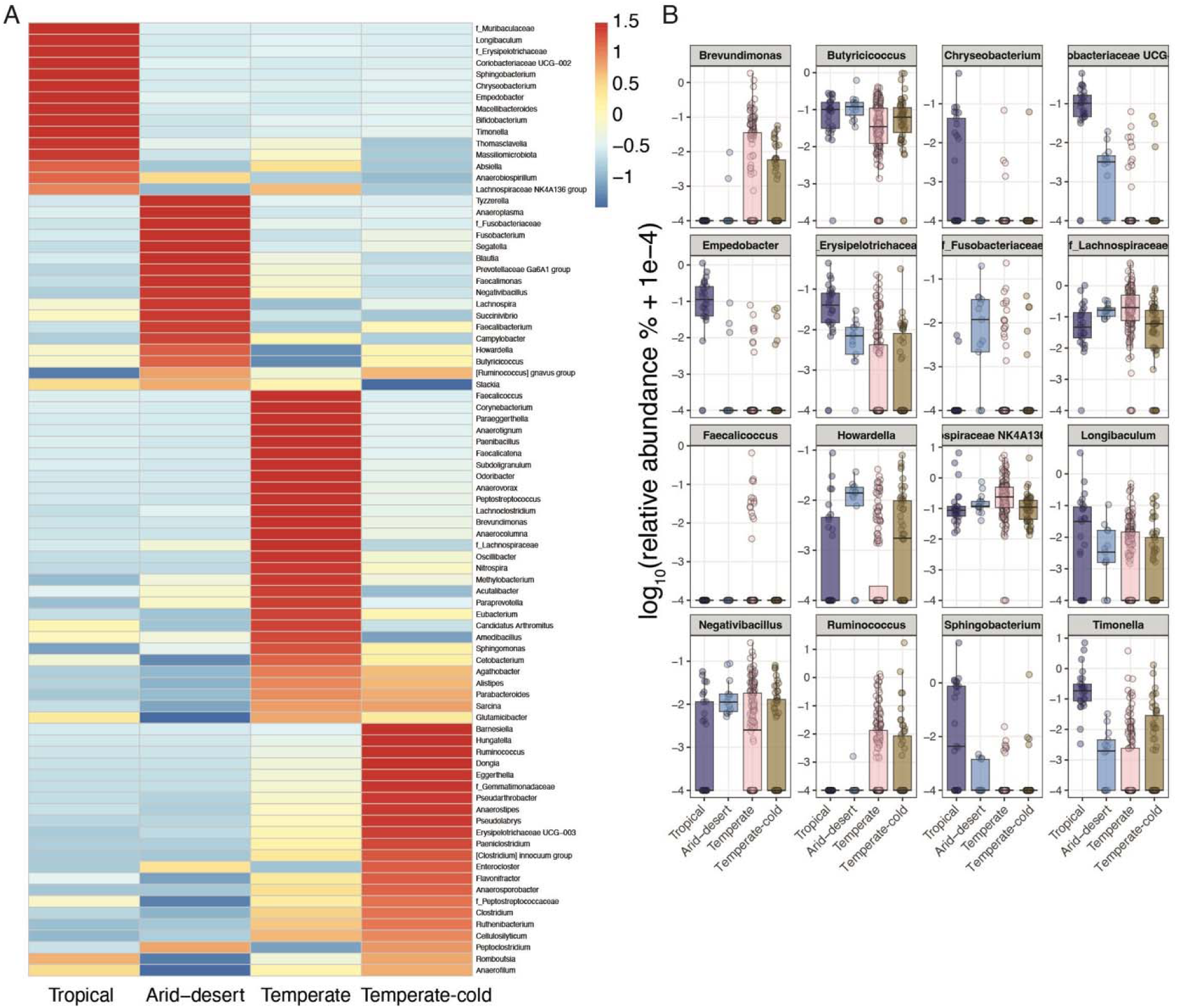
Climate-associated genera show zone-specific abundance structure. **(A)** Heatmap of mean standardized abundance (z scores) for genera showing significant large-effect associations with climate category in the multivariable MaAsLin2 model. Genera were identified using an FDR-adjusted q value < 0.05 and an absolute coefficient > 1. Columns represent climate categories, and rows are ordered by the climate category in which each genus shows its highest mean abundance. In contrast to lifestyle-associated genera, climate-associated genera form discrete climate-enriched blocks rather than a directional gradient. **(B)** Boxplots showing the subset of climate-associated genera that remained associated with climate category after excluding genera that also associated with lifestyle or age. Points represent individual dogs; boxes show the median and interquartile range. Values are plotted as log10(relative abundance + 1e-4). Several of these climate-specific genera, including *Sphingobacterium*, *Chryseobacterium*, and *Empedobacter*, were most enriched in tropical environments, consistent with climate-linked environmental exposure signals.

**Figure S9.**
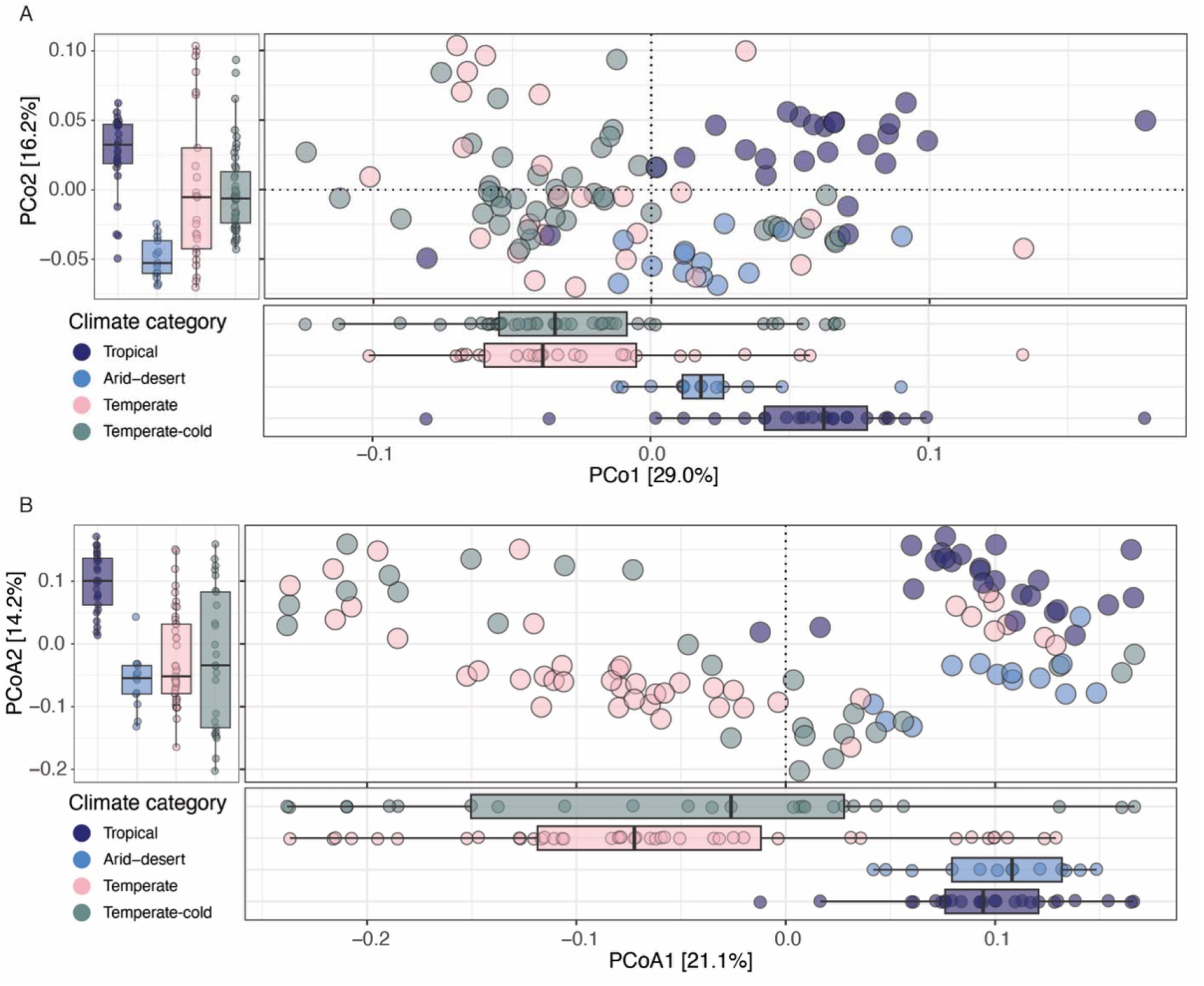
Climate-associated microbiome differences remain detectable within industrialized dogs. Principal coordinate analysis (PCoA) of genus-level gut microbiome composition among industrialized dogs sampled across four climate categories: tropical, arid-desert, temperate, and temperate-cold. **(A)** Weighted UniFrac PCoA. **(B)** Bray-Curtis PCoA. Points represent individual dogs and are colored by climate category. Marginal boxplots summarize sample distributions along the first two ordination axes. In both distance frameworks, industrialized dogs from different climate categories showed significant compositional differences (PERMANOVA, P < 0.05), indicating that climate- and geography-associated structure remains detectable even within a shared industrialized lifestyle background.

**Figure S10.**
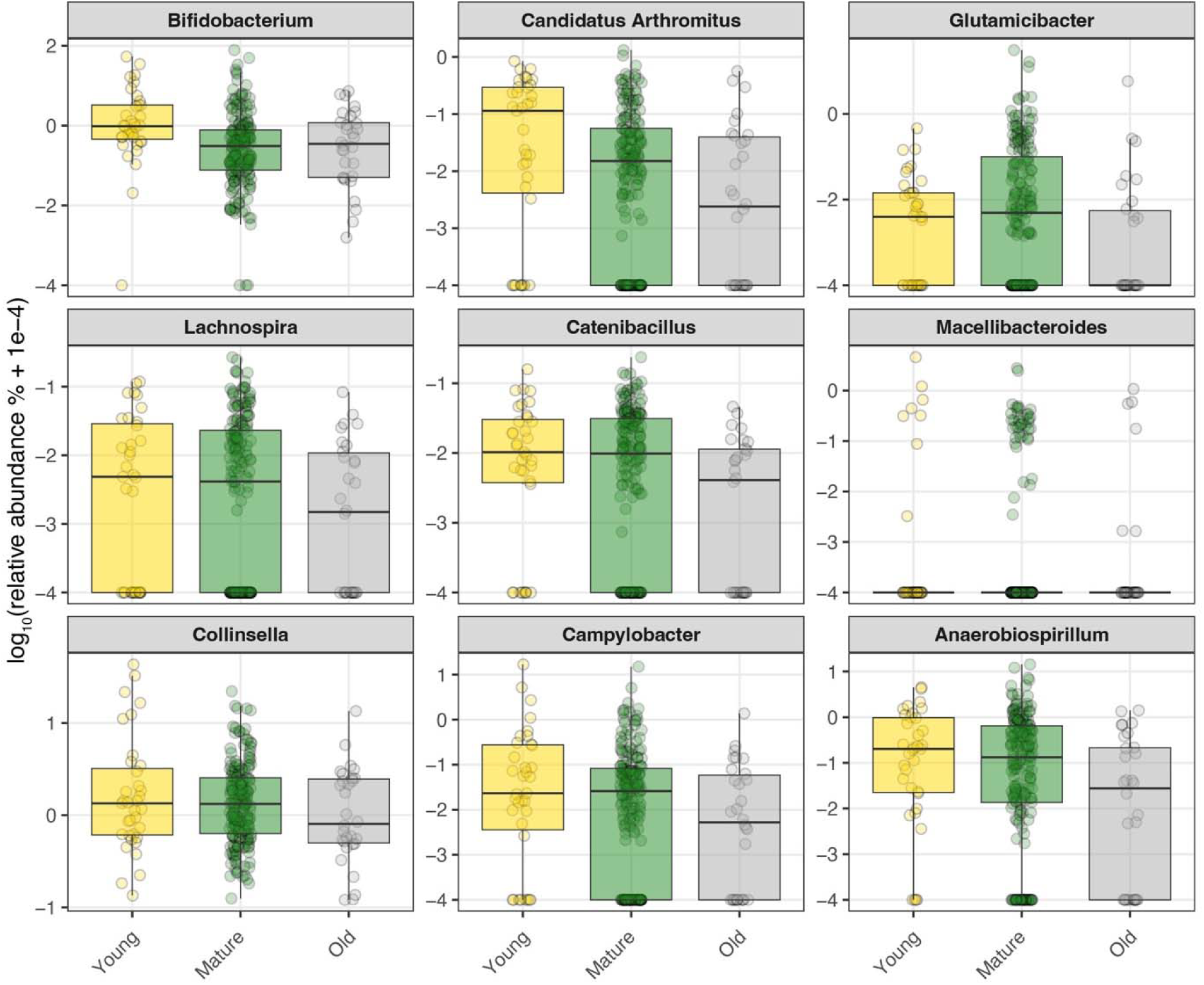
Age-associated genera identified in the multivariable MaAsLin2 model. Boxplots show log-transformed relative abundances of the nine genera with significant large-effect associations with age group in the multivariable model (FDR-adjusted q < 0.05 and absolute MaAsLin2 coefficient > 1). Points represent individual dogs and are colored by age group (young, mature, and old). Among these taxa, *Bifidobacterium* showed a decline with age, consistent with microbiome maturation, whereas the remaining age-associated genera showed comparatively modest differences across age groups.

**Figure S11.**
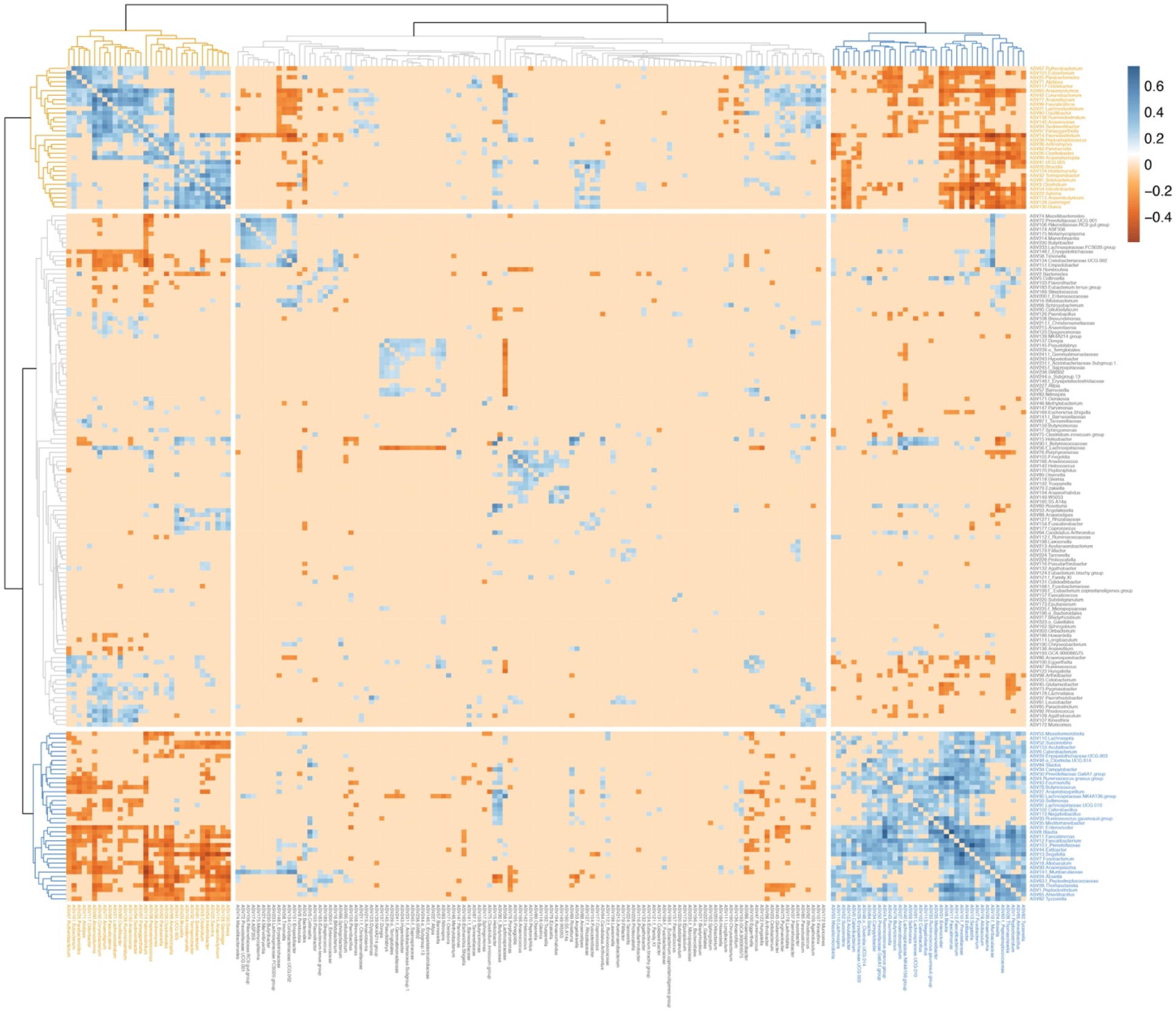
Genus-level co-abundance structure among prevalence-shifting canine gut microbiome taxa. Heatmap of pairwise genus-genus associations inferred using FastSpar among genera included in the co-abundance group analysis. Rows and columns represent genera, ordered by hierarchical clustering of the filtered correlation matrix. Colors indicate FastSpar correlation coefficients, with blue representing positive associations and orange representing negative associations. Only associations passing the filtering threshold used for network construction are emphasized (absolute rho > 0.2 and FDR-adjusted P < 0.05). The dendrograms show clustering of genera into co-abundance groups (CAGs), which resolved three major modules: two tightly connected CAGs and one more diffuse cluster. These CAG assignments were used for the network visualization and lifestyle-level CAG abundance summaries shown in Fig. 4B and C.

**Figure S12.**
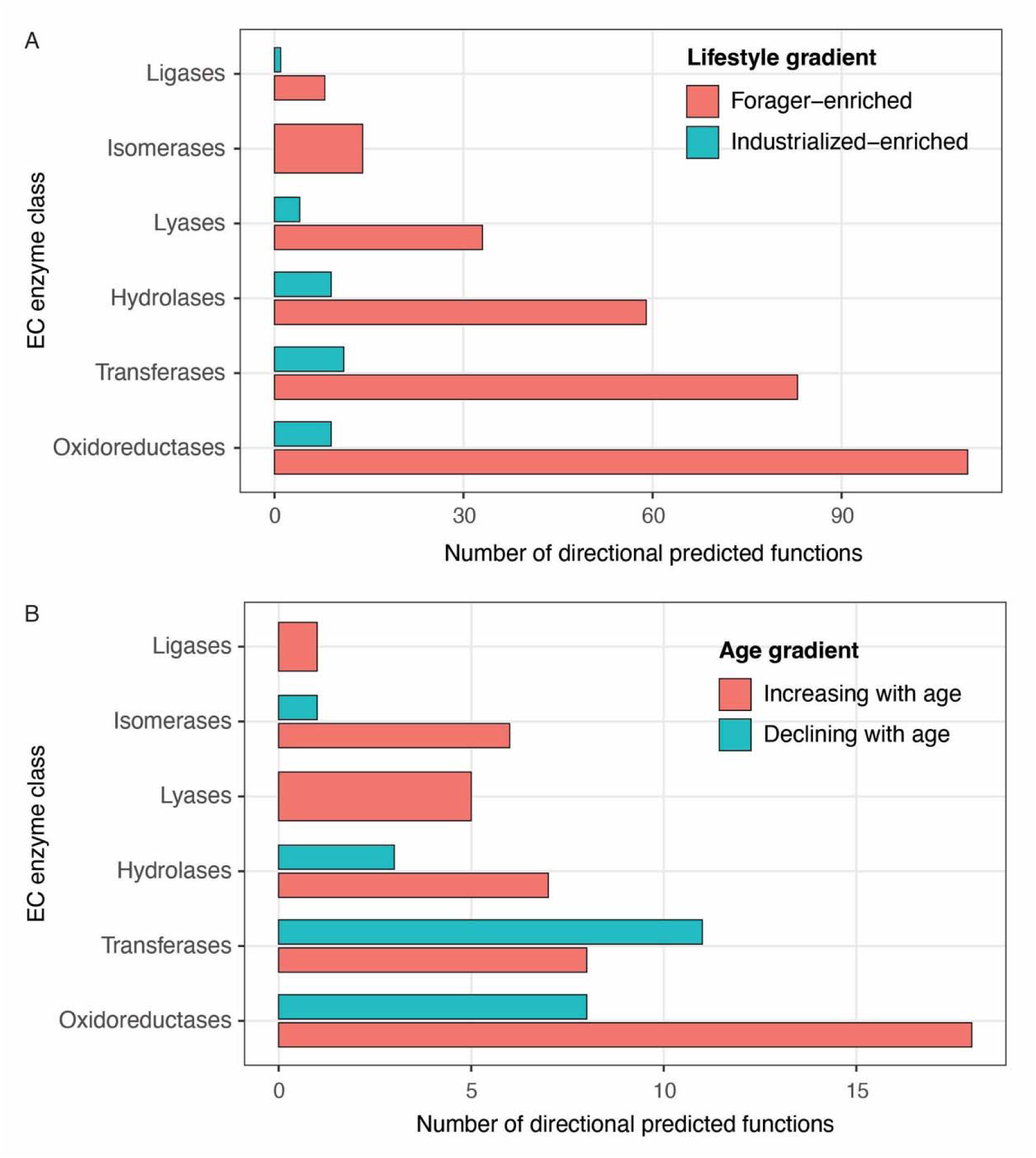
Predicted enzyme functions associated with lifestyle and age gradients. Distribution of directional predicted enzyme functions across broad EC enzyme classes. Predicted functional profiles were inferred from 16S rRNA gene data using PICRUSt2 and analyzed with MaAsLin2 models including lifestyle, climate category, and age group. **(A)** Predicted enzyme functions showing monotonic shifts across the forager-agrarian-urban-industrialized lifestyle gradient, classified as forager-enriched or industrialized-enriched according to endpoint direction. **(B)** Predicted enzyme functions showing directional associations with age group, classified as increasing or declining with age. Bars indicate the number of significant directional predicted functions in each EC enzyme class. Predicted functions should be interpreted as inferred functional potential rather than directly measured metagenomic activity.

**Figure S13.**
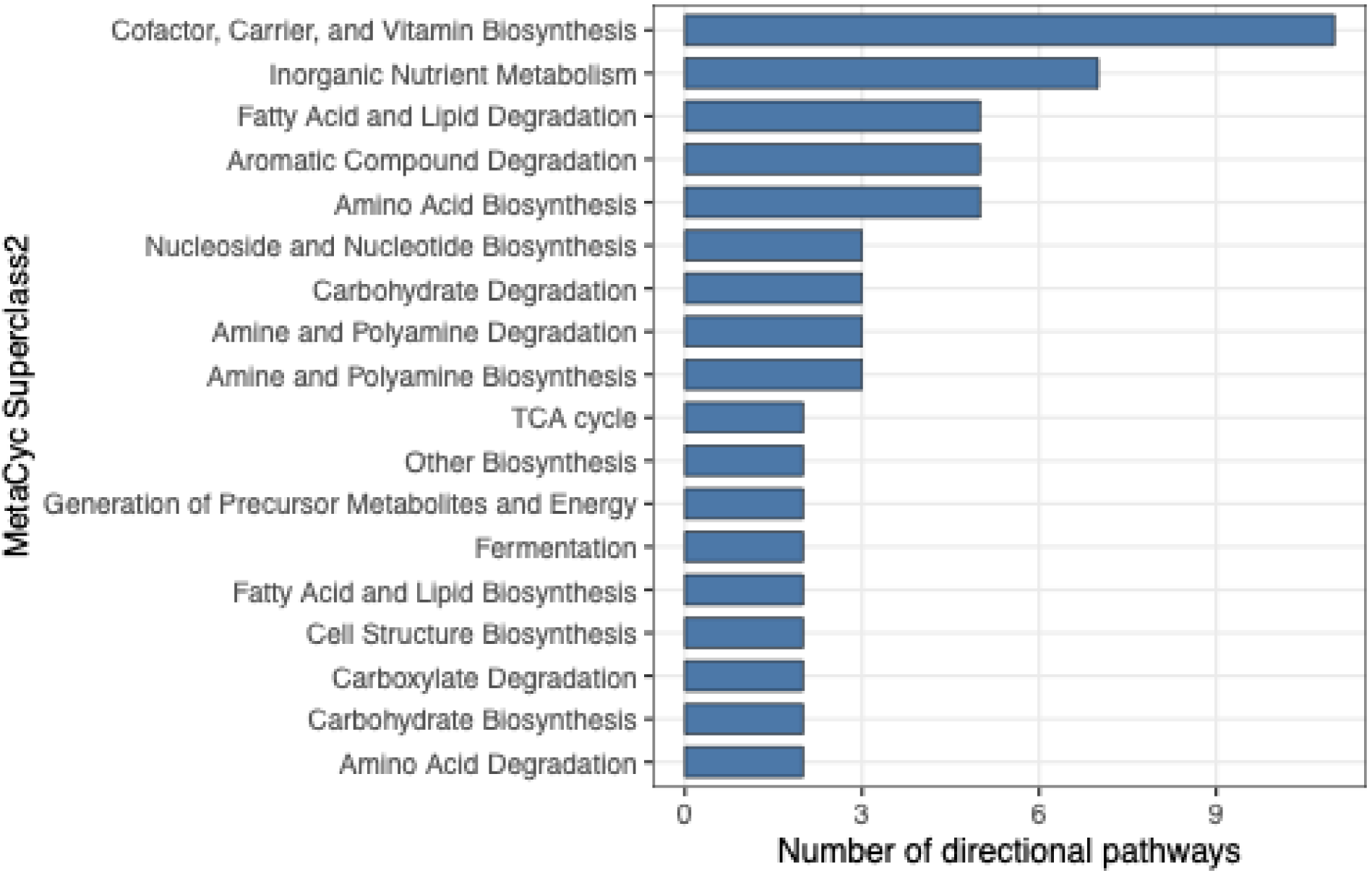
Predicted pathway profiles shift across the canine lifestyle gradient. Bar plot showing the distribution of the 85 lifestyle-associated directional predicted MetaCyc pathways across MetaCyc superclass categories. Pathways were identified using the same monotonic lifestyle-gradient criterion applied to genus-level taxa across forager, agrarian, urban, and industrialized dogs. Categories with fewer than two directional pathways were omitted for visualization. Because these pathway profiles were inferred from 16S rRNA gene data, they should be interpreted as predicted functional potential rather than directly measured metagenomic activity.

**Figure S14.**
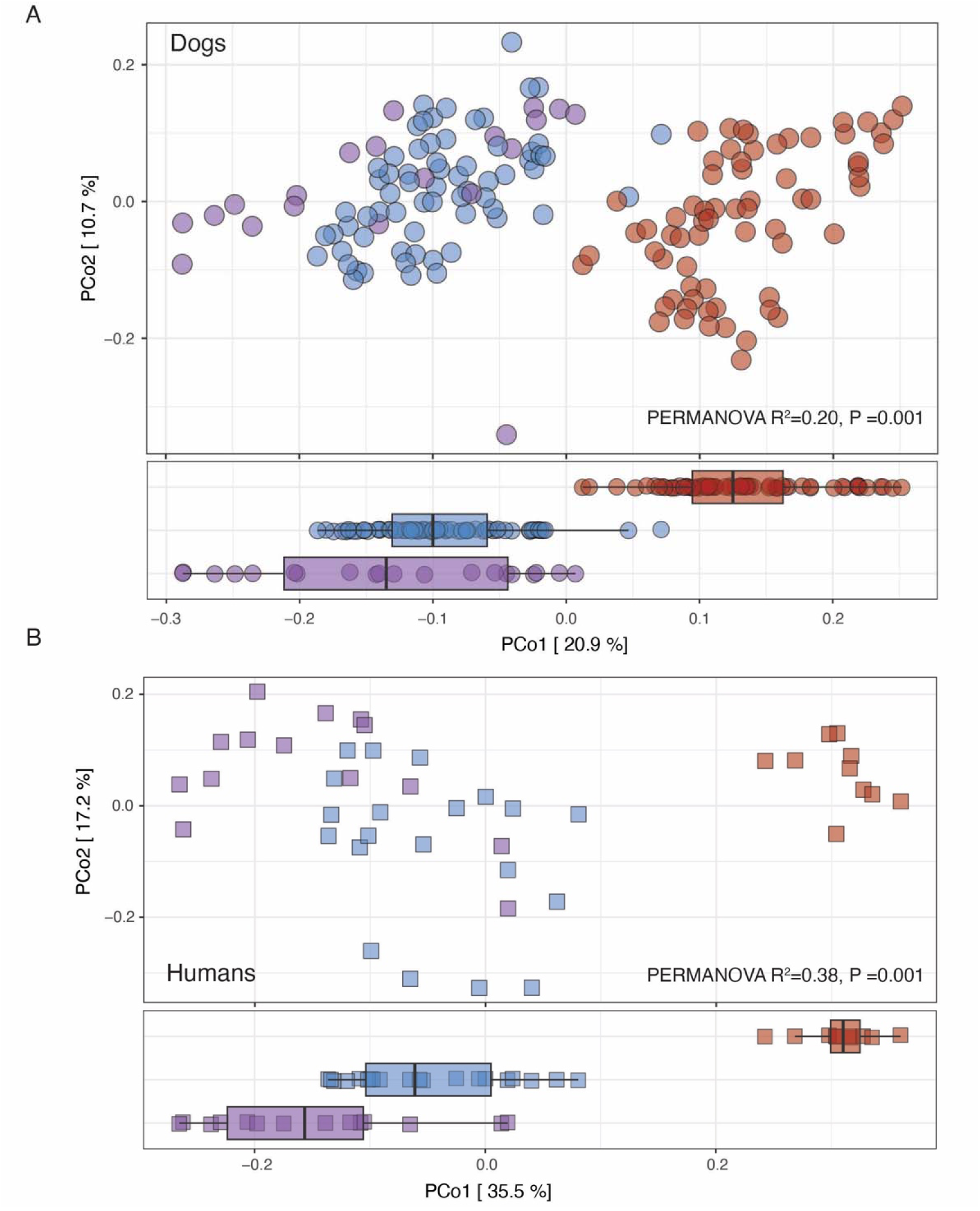
Host-specific lifestyle-associated microbiome structure in dogs and humans. Separate Bray-Curtis PCoA analyses of genus-level microbiome profiles from dogs and humans sampled across matched forager, agrarian, and industrialized lifestyle categories. **(A)** Dog samples from the matched cross-host comparison. **(B)** Human samples from Jha et al. 2018 reprocessed using the same taxonomic workflow and restricted to genus-level profiles. Points represent individual samples and are colored by lifestyle category; marginal boxplots show distributions along PCo1. In both hosts, industrialized samples separate from forager and agrarian samples along the leading ordination axis, indicating lifestyle-associated microbiome restructuring within each host. PERMANOVA R^2^ and P values indicate the association between lifestyle and Bray-Curtis community composition.

**Figure S15.**
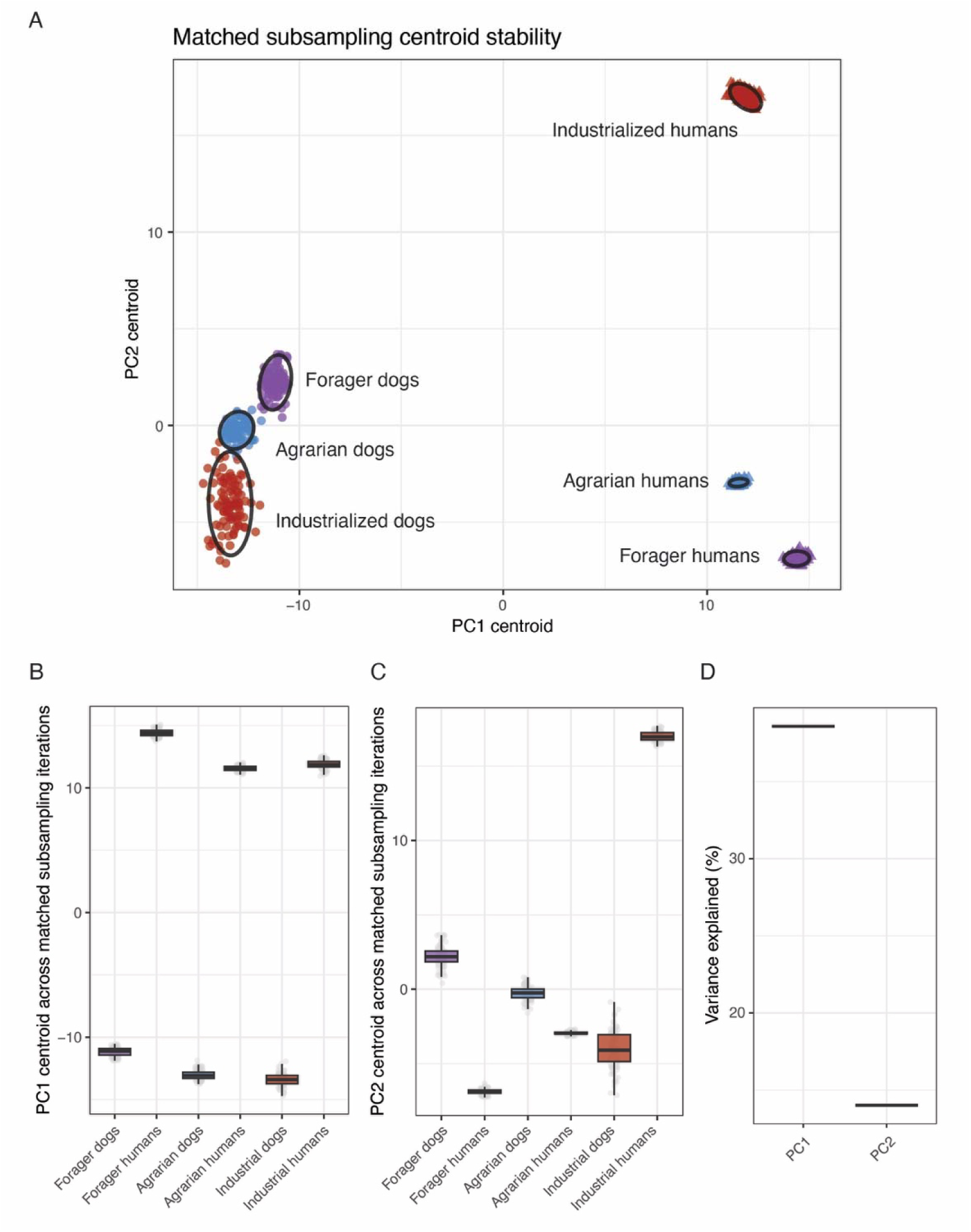
Robustness of dog–human PCA patterns to matched dog subsampling. Dogs were repeatedly subsampled within lifestyle categories to match the number of human samples: foragers, n = 14; agrarian, n = 20; industrialized, n = 10. In each of 100 iterations, the matched dog subset was merged with the full human dataset using shared genera, and PCA was repeated using the same genus-level abundance transformation as in the primary analysis. **A**, Host × Lifestyle centroid positions across iterations. Points represent centroids from individual subsampling iterations; ellipses summarize centroid dispersion. **B–C**, distributions of PC1 and PC2 centroid coordinates across iterations. **D**, variance explained by PC1 and PC2 across iterations. Stable centroid positions indicate that the observed dog–human PCA structure is robust to matched lifestyle-stratified dog subsampling.

**Figure S16.**
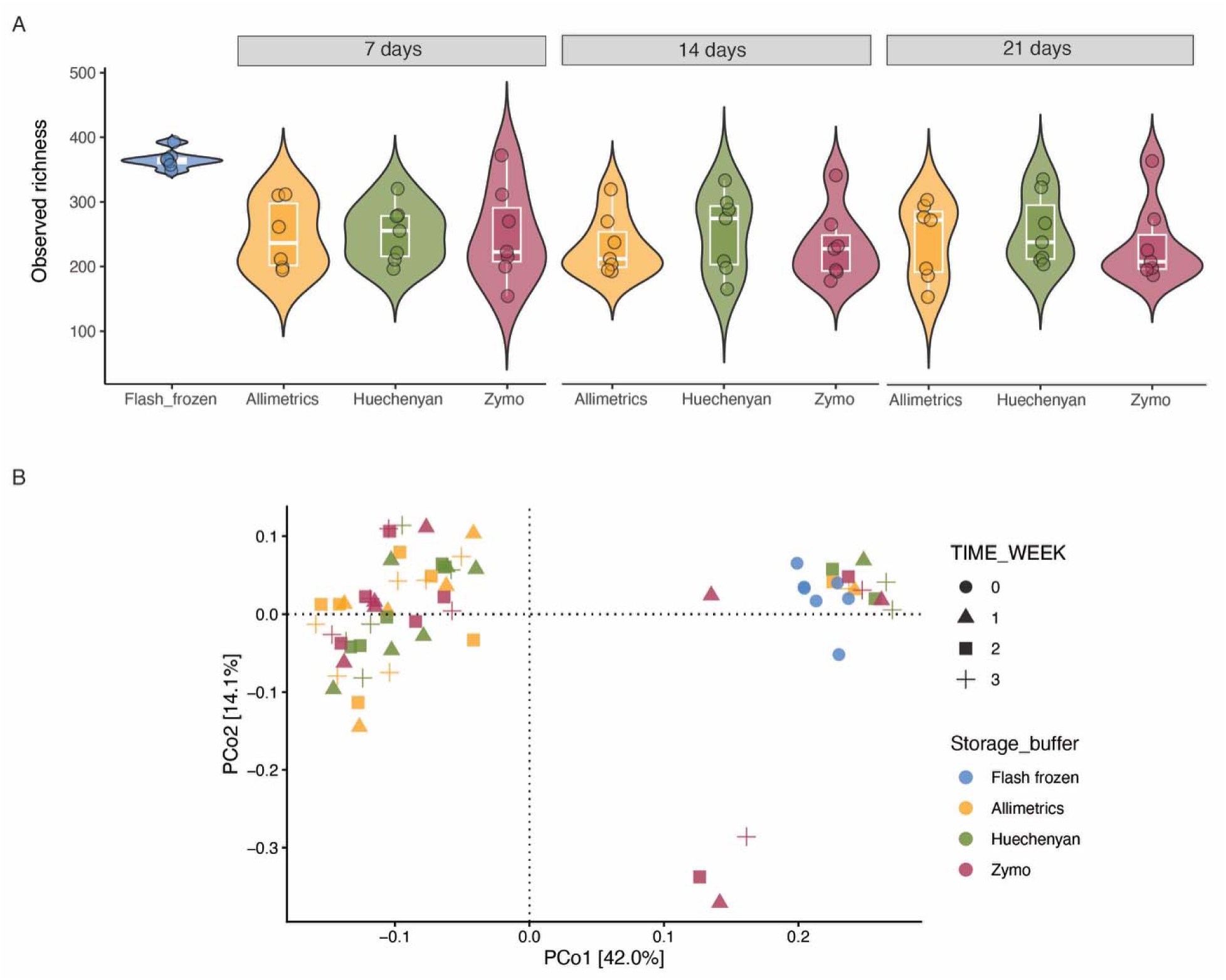
Optimization of field-compatible fecal sample preservation methods. Fecal samples from seven pet dogs at New York University Abu Dhabi were used to compare flash freezing with three commercial preservation systems: Allimetrics fecal collection tubes, Huechenyan fecal collection tubes, and Zymo DNA/RNA Shield Fecal Collection Tubes. Fresh fecal samples were transported to the laboratory on dry ice within 30 minutes, divided into aliquots, and either flash frozen at −80°C or stored in preservation buffer at room temperature for 7, 14, or 21 days before DNA extraction. **(A)** Observed richness differed between flash-frozen and buffer-preserved samples, with higher richness in flash-frozen aliquots, but did not differ significantly among preservation buffers or across room-temperature storage durations. **(B)** Bray-Curtis PCoA showed a significant effect of preservation method on overall microbiota composition (PERMANOVA R² = 0.14, P = 0.0004). Pairwise PERMANOVA indicated that flash-frozen samples differed from buffer-preserved samples, whereas microbial profiles did not differ significantly among the three preservation buffers or among buffer-preserved samples stored for 7, 14, or 21 days. Because the preservation buffers performed comparably under field-relevant storage conditions, the cost-effective Huechenyan kit was selected for large-scale field sampling.

**Figure S17.**
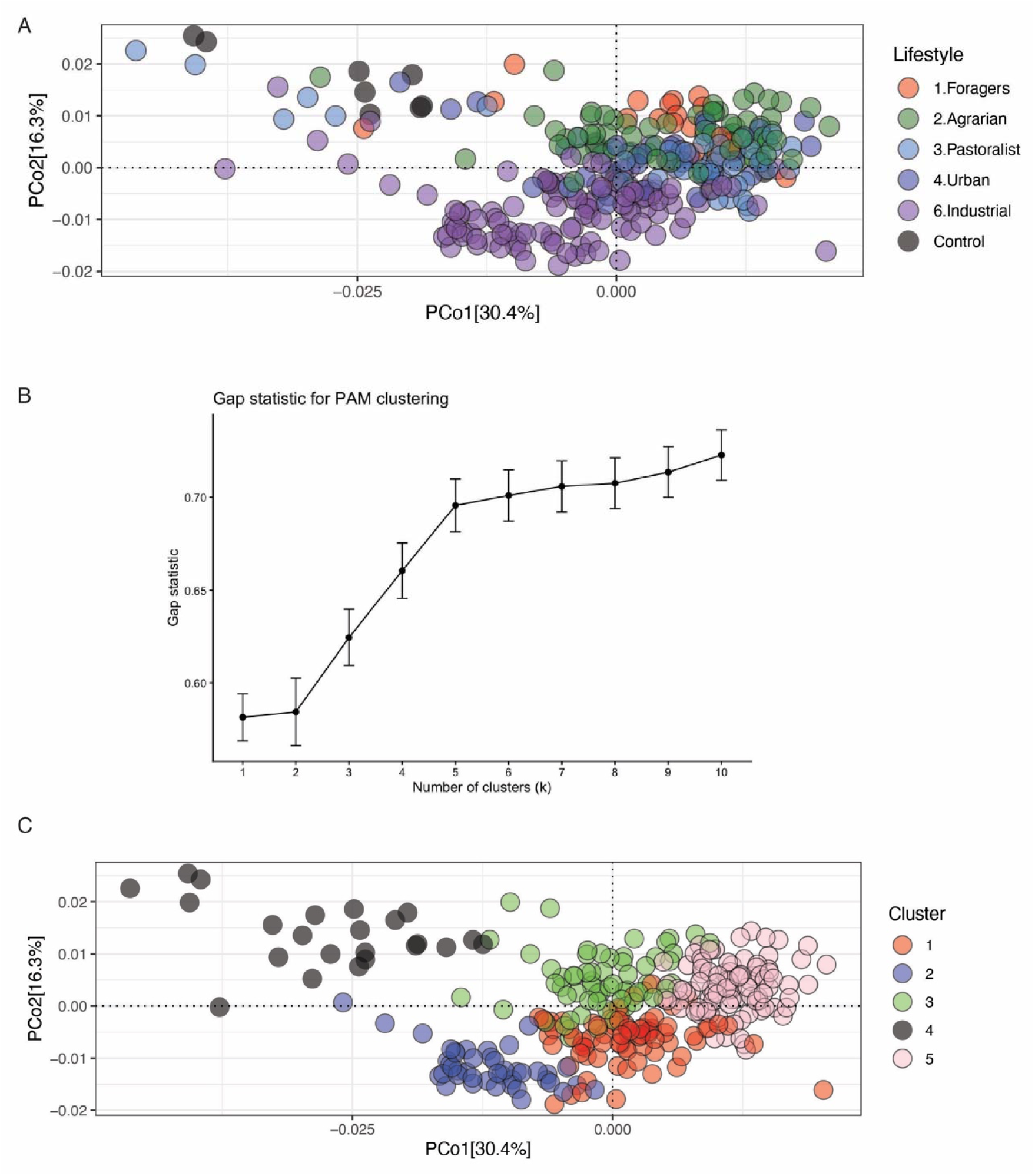
Identification of contamination-like samples using extraction controls. **(A)** Principal coordinate analysis (PCoA) of weighted UniFrac distances calculated from genus-level abundance profiles for 286 fecal samples and 13 negative extraction controls. Points represent samples colored by lifestyle category or control status. Negative extraction controls clustered separately from most fecal samples. **(B)** Gap statistic for partitioning around medoids (PAM) clustering across k = 1-10. The gap statistic increased up to k = 5, after which additional clusters produced smaller gains. **(C)** PCoA of the same weighted UniFrac distances colored by the k = 5 PAM cluster assignment. At k = 5, an outlier control-enriched cluster (gray) contained 8 of 13 retained negative extraction controls and 15 fecal samples that clustered with controls; these 15 fecal samples were excluded from downstream analyses.

### Additional File 2. Supplementary Tables

**Table S1. Metadata for domestic dog fecal samples included in the study.** Sample-level metadata for 261 domestic dogs sampled across Nepal, Thailand, the United Arab Emirates, and the United States. The table includes sample identifiers, country, sampling location, geographic coordinates, altitude, climate category, lifestyle classification, diet category, shelter status, age category, sex, health status, disease status, and recent antibiotic exposure. Four dogs with reported recent antibiotic exposure were excluded from downstream microbiome analyses, yielding 257 dogs for the primary analyses.

**Table S2. Associations between canine gut microbiome composition and measured host or ecological variables.**

PERMANOVA and EnvFit results for genus-level weighted UniFrac and Bray-Curtis distance matrices. The upper section shows analyses performed on the full dataset of 257 dogs after excluding individuals with recent antibiotic exposure. Marginal PERMANOVA models included lifestyle, climate category, altitude category, shelter status, and age group, with effects estimated using by = "margin". EnvFit analyses assessed alignment of each variable with the first three ordination axes. The lower section shows sensitivity analyses performed on a restricted subset of 107 mature, non-shelter dogs sampled from temperate climate regions, excluding Thai and UAE dogs. In this restricted dataset, lifestyle remained strongly associated with microbiome composition in both weighted UniFrac and Bray-Curtis analyses. P values were calculated using 999 permutations.

**Table S3. Associations between canine gut microbiome alpha diversity and measured host or ecological variables.** Gamma generalized linear model results testing associations between sample-level alpha diversity and lifestyle, climate category, shelter status, and age group in the 257-dog analysis dataset after excluding dogs with recent antibiotic exposure. Alpha diversity was measured as observed ASV richness, Shannon diversity, and Faith’s phylogenetic diversity after rarefaction to 5,000 reads per sample. Industrialized lifestyle, tropical climate, non-shelter status, and young age group were used as reference categories. Estimates, standard errors, t-values, nominal P values, and Benjamini-Hochberg-adjusted q values are shown. Q values were calculated across all non-intercept coefficients from the three alpha-diversity models. No predictor remained significant after multiple-testing correction, indicating that lifestyle-associated variation was reflected primarily in community composition rather than overall within-sample diversity.

**Table S4. Lifestyle-associated genus-level differential abundance in the canine gut microbiome.** MaAsLin2 results from the lifestyle-only differential abundance model testing genus-level associations with canine lifestyle category in the 257-dog analysis dataset after excluding dogs with recent antibiotic exposure. Industrialized dogs were used as the reference group, and coefficients indicate the direction and magnitude of association for each non-industrialized lifestyle category relative to industrialized dogs. The table includes genus-level taxonomy, metadata variable, lifestyle contrast, MaAsLin2 coefficient, standard error, sample number, number of non-zero samples, nominal P value, Benjamini-Hochberg-adjusted q value, and absolute coefficient. Multiple testing correction was performed using the Benjamini-Hochberg procedure. Genera were considered significant large-effect associations if q < 0.05 and absolute coefficient > 1.

**Table S5. Directional genus-level shifts across the canine lifestyle gradient.** Subset of lifestyle-associated genera from Table S4 that showed monotonic abundance patterns across the forager-agrarian-urban-industrialized lifestyle axis. Pastoralist dogs were excluded from this gradient analysis because they represented a distinct high-altitude ecological context rather than an intermediate stage along the forager-to-industrialized continuum. MaAsLin2 coefficients are shown for urban, agrarian, and forager dogs relative to industrialized dogs, which served as the reference group. Genera were retained if the forager-versus-industrialized endpoint contrast met the large-effect threshold defined in Table S4 (FDR-adjusted q < 0.05 and absolute coefficient > 1) and if coefficients followed a monotonic increase or decrease across the lifestyle axis.

**Table S6. Multivariable genus-level associations with lifestyle, climate category, and age group.** MaAsLin2 results from the multivariable differential abundance model testing associations between genus-level relative abundance and lifestyle, climate category, and age group in the 257-dog analysis dataset after excluding dogs with recent antibiotic exposure. Industrialized dogs, tropical climate, and young dogs were used as the reference categories. The table includes genus-level taxonomy, metadata variable, contrast level, MaAsLin2 coefficient, standard error, sample number, number of non-zero samples, nominal P value, Benjamini-Hochberg-adjusted q value, and absolute coefficient. This model was used to assess whether lifestyle-associated genera remained associated with lifestyle after adjustment for climate category and age group.

**Table S7. Prevalence-defined VANISH-like and BloSSUM-like genera in the canine gut microbiome.** Genera showing large prevalence differences between non-industrialized and industrialized dogs. Forager and agrarian dogs were grouped as the non-industrialized comparison set and compared with industrialized dogs using genus-level presence-absence profiles. Genera were classified as VANISH-like if they were significantly more prevalent in forager/agrarian dogs and as BloSSUM-like if they were significantly more prevalent in industrialized dogs. Genera were retained if they showed an absolute prevalence difference ≥ 30% and a Benjamini-Hochberg-adjusted Fisher’s exact test P value < 0.05. The table includes genus-level taxonomy, presence and absence counts in forager/agrarian and industrialized dogs, prevalence proportions, prevalence difference, Fisher’s exact test estimate, nominal P value, and Benjamini-Hochberg-adjusted P value.

**Table S8. Directional shifts in predicted enzyme functions across the canine lifestyle gradient.** Predicted enzyme functions showing monotonic abundance patterns across the forager-agrarian-urban-industrialized lifestyle axis. Functional profiles were inferred from 16S rRNA gene data using PICRUSt2 and analyzed with MaAsLin2 models including lifestyle, climate category, and age group. Industrialized dogs were used as the reference group, and pastoralist dogs were excluded from the monotonic-gradient analysis because they represented a distinct high-altitude ecological context rather than an intermediate stage along the forager-to-industrialized continuum. Predicted enzyme functions were retained if the forager-versus-industrialized endpoint contrast showed a significant large-effect association, defined as FDR-adjusted q < 0.05 and absolute MaAsLin2 coefficient > 1, and if coefficients followed a monotonic increase or decrease across the lifestyle axis.

**Table S9. Directional shifts in predicted MetaCyc pathways across the canine lifestyle gradient.** Predicted MetaCyc pathways showing monotonic abundance patterns across the forager-agrarian-urban-industrialized lifestyle axis. Pathway profiles were inferred from 16S rRNA gene data using PICRUSt2 and analyzed with MaAsLin2 models including lifestyle, climate category, and age group. Industrialized dogs were used as the reference group, and pastoralist dogs were excluded from the monotonic-gradient analysis because they represented a distinct high-altitude ecological context rather than an intermediate stage along the forager-to-industrialized continuum. Predicted pathways were retained if the forager-versus-industrialized endpoint contrast showed a significant large-effect association, defined as FDR-adjusted q < 0.05 and absolute MaAsLin2 coefficient > 1, and if coefficients followed a monotonic trend across the lifestyle axis. Pathway superclass annotations were obtained from MetaCyc using pathway identifiers.

**Table S10. Multivariable associations between predicted MetaCyc pathways and host or ecological variables.** MaAsLin2 results from the full multivariable model testing associations between predicted MetaCyc pathway abundance and lifestyle, climate category, and age group in the 257-dog analysis dataset after excluding dogs with recent antibiotic exposure. Pathway profiles were inferred from 16S rRNA gene data using PICRUSt2 and should be interpreted as predicted functional potential rather than directly measured metagenomic activity. Industrialized dogs, tropical climate, and young dogs were used as reference categories. The table includes MetaCyc pathway identifiers, pathway superclass annotations, pathway names, metadata variable, contrast level, MaAsLin2 coefficient, standard error, sample number, number of non-zero samples, nominal P value, Benjamini-Hochberg-adjusted q value, and PICRUSt2 feature identifier. This table contains all significant pathway-level contrasts from the multivariable model, including the lifestyle-gradient pathways summarized separately in Table S9, as well as additional lifestyle-, climate-, and age-associated pathway contrasts that did not meet the monotonic lifestyle-gradient criterion.

**Table S11. Shared genus-level taxa and lifestyle associations in dogs and humans.** Genera shared between the dog and human datasets and MaAsLin2 results from host-specific lifestyle-gradient analyses. Dog and human datasets were restricted to the 98 genera detected in both hosts and analyzed separately across matched forager, agrarian, and industrialized lifestyle categories. Industrialized samples were used as the reference group within each host. The table includes genus taxonomy, host specfic MaAsLin2 coefficients, standard errors, nominal P values, Benjamini-Hochberg-adjusted q values, monotonic gradient direction, endpoint effect size, and indicators of whether each genus showed same-direction or opposite-direction lifestyle-associated responses across hosts. NA indicates that a monotonic lifestyle gradient was not assigned in one or both hosts. This table was used to assess whether the same genus-level taxa responded similarly to lifestyle transitions in dogs and humans.

## Notes

### Competing Interest Statement

The authors have declared no competing interest.

